# HNF4A maintains proximal tubule identity and limits injury-associated cell states in the adult mouse kidney

**DOI:** 10.64898/2026.09.10.750757

**Authors:** Fariba Nosrati, Zeinab Dehghani-Ghobadi, Eunah Chung, Christopher Ahn, Hee-Woong Lim, Joo-Seop Park

## Abstract

**Introduction:** HNF4A is required for proximal tubule maturation during kidney development, but its role in maintaining proximal tubule identity in the adult kidney has not been defined. Following acute kidney injury, *Hnf4a* expression is rapidly suppressed in proximal tubule cells, coinciding with loss of mature epithelial features and activation of injury-associated transcriptional programs. It remains unknown whether HNF4A loss is sufficient to drive injury-associated transcriptional changes.

**Methods:** We conditionally deleted *Hnf4a* in mature proximal tubules using *Slc34a1Cre* in mice and analyzed the consequences of *Hnf4a* loss in adult kidneys. To identify genes directly regulated by HNF4A, we performed proximal tubule-specific transcriptional profiling together with genome-wide mapping of HNF4A binding sites using CUT&RUN.

**Results:** In *Hnf4a* mutant kidneys, HNF4A protein initially persisted in proximal tubules but was progressively lost from convoluted proximal tubule cells. Loss of *Hnf4a* resulted in downregulation of proximal tubule-specific transport and metabolic genes, accompanied by reactivation of developmental and injury-associated genes. CUT&RUN analysis revealed that HNF4A directly regulates genes predominantly involved in solute transport and metabolic processes.

**Conclusions:** These findings identify HNF4A as a key regulator of proximal tubule identity and homeostasis in the adult mouse kidney. Genetic loss of *Hnf4a* in mature proximal tubules leads to loss of mature proximal tubule gene expression and activation of injury-associated transcriptional programs, implicating *Hnf4a* suppression as a potential contributor to maladaptive repair after kidney injury.

**Translational Statement:** Proximal tubule dysfunction is a major contributor to acute kidney injury and progression to chronic kidney disease. Here, we show that HNF4A is required to maintain mature proximal tubule identity in the adult kidney. Loss of HNF4A suppresses transport and metabolic programs and induces transcriptional features associated with injured, failed-repair, and immature proximal tubule states, even in the absence of exogenous kidney injury. These findings suggest that suppression of HNF4A-dependent transcriptional programs may contribute to maladaptive proximal tubule remodeling after injury and identify HNF4A-regulated pathways as potential mechanisms for preserving differentiated proximal tubule function.

## Introduction

The mammalian kidney maintains systemic homeostasis by filtering blood to remove metabolic waste while selectively reabsorbing and secreting solutes along specialized nephron segments. The proximal tubule (PT), the largest segment of the nephron, reabsorbs most filtered solutes and water and plays central roles in renal metabolism.^1–3^ Because of its high metabolic demand and exposure to filtered toxins, the PT is particularly susceptible to injury.^4–6^ Consequently, PT cell dysfunction or loss is a major contributor to acute kidney injury (AKI) and to the progression from AKI to chronic kidney disease (CKD).^7–12^ Proper differentiation and maintenance of PT cells are therefore essential for preserving renal function and tissue homeostasis.^13^

HNF4A is a transcription factor that is critical for epithelial differentiation and organ function in multiple tissues, including the kidney.^14–17^ During nephrogenesis, *Hnf4a* is expressed specifically in both immature and mature PT cells.^15, 16, 18^ Our previous studies demonstrated that immature PT cells can form following *Hnf4a* deletion, but they fail to differentiate into mature PT cells, indicating that HNF4A is dispensable for PT specification but required for PT maturation.^15, 16^ In these studies, *Hnf4a* was conditionally deleted using either Six2TGC (the *Six2* transgenic Cre), which targets nephron progenitors, or Osr2Cre, which targets epithelial progenitors in the S-shaped body.^19–21^ Because both Cre drivers are active before the onset of *Hnf4a* expression in the developing kidney, mature PT cells fail to form in these mutants. As a result, these models delete *Hnf4a* prior to, rather than after, PT maturation, and therefore cannot address whether HNF4A is required to maintain PT identity in the adult kidney.

In addition to its developmental role, recent studies have suggested that HNF4A may play an important role in the response of PT cells to injury.^22^ Injured PT cells downregulate mature transport and metabolic programs while activating inflammatory, regenerative, and repair-associated pathways.^23, 24^ Notably, *Hnf4a* expression is reduced in injured, dedifferentiated, and failed-repair PT cells,^7, 25–28^ consistent with loss of mature PT identity during AKI and CKD progression. These observations raise the possibility that suppression of *Hnf4a* is not merely a consequence of PT injury but may actively contribute to injury-associated transcriptional changes. However, whether loss of HNF4A function in mature PT cells is sufficient to drive these changes remains unknown.

In the present study, we deleted *Hnf4a* specifically in differentiated PT cells using Slc34a1Cre,^18^ thereby bypassing the developmental defects observed in earlier models and enabling direct investigation of HNF4A function in mature PT cells. Using a combination of histologic, transcriptomic, and HNF4A chromatin occupancy analyses, we determined how HNF4A maintains PT identity and whether loss of HNF4A is sufficient to induce injury-associated transcriptional programs in the adult kidney in the absence of exogenous injury.

## Methods

### Mice

All mouse alleles used in this study have been previously published: *Rosa26^NuTRAP^* (JAX:029899),^29^ *Hnf4a^c^* (JAX:004665),^30^ and *Slc34a1^Cre^*(JAX:040319).^18^ Animals were housed in a controlled facility under a 12-hour light/12-hour dark cycle with ad libitum access to a standard chow diet and water. Both male and female mice were used in this study unless otherwise noted. All animal handling and experimental procedures were performed in accordance with the National Institutes of Health Guide for the Care and Use of Laboratory Animals. The protocols were reviewed and approved by the Institutional Animal Care and Use Committee of Northwestern University.

### Immunofluorescence staining and imaging

Kidneys were fixed in 4% paraformaldehyde in phosphate-buffered saline (PBS) for 20 minutes, followed by overnight incubation in 10% sucrose in PBS at 4°C. Tissues were embedded in Optimal Cutting Temperature compound (Tissue-Tek O.C.T. Compound, Sakura Finetek USA 4583). Cryosections (8-10 μm) were incubated overnight with primary antibodies (Supplemental Table 1) in PBS containing 5% heat-inactivated sheep serum and 0.1% Triton X-100. Fluorophore-labeled secondary antibodies were used for indirect detection (Supplemental Table 1). Images were acquired using a Nikon Ti-2 inverted widefield microscope equipped with an Orca Fusion camera and a Lumencor Spectra III light source. Images shown in the figures are representative fields from analyses performed on at least four independent control mice and four independent *Hnf4a* mutant mice per staining condition.

### Fluorescence-activated cell sorting and bulk RNA-seq

Kidneys from *Hnf4a* mutant and control mice at postnatal day 61 (P61), carrying the Slc34a1Cre-driven Rosa26-NuTRAP reporter, were dissociated using TrypLE Select Enzyme (Thermo A1217701) with repeated pipetting. Cells were resuspended in PBS containing 1% FBS and 10 mM EDTA, then filtered through a 40 µm nylon strainer (BD Falcon 352340). GFP-positive cells were isolated using BD FACSAria at the Robert H. Lurie Comprehensive Cancer Center Flow Cytometry Core Facility at Northwestern University. Total RNA was extracted using the Single Cell RNA Purification Kit (Norgen Biotek 51800), and mRNA was isolated with the NEBNext Poly(A) Magnetic Isolation Module (E7490L, New England Biolabs). Subsequent steps were done as described previously.^31^ Libraries were sequenced on Illumina NovaSeq X Plus or Element AVITI at the NUSeq Core facility at Northwestern University. Bulk RNA-seq was performed on GFP-positive proximal tubule cells isolated from four control mice (two males and two females) and four *Hnf4a* mutant mice (two males and two females), with each mouse serving as an independent biological replicate.

### PT morphometric analysis

For PT morphometric analysis, kidney sections were stained for GFP, laminin, and Hoechst. GFP+ regions were used to define PT areas, and the resulting GFP mask was applied to the laminin and Hoechst channels to restrict the analysis to GFP+ PTs. Laminin+ basement membranes were used to delineate individual PT profiles. Convoluted and straight PTs were distinguished based on anatomical location and morphology. Tubular diameter was measured in Fiji/ImageJ as the shortest distance between opposing laminin+ basement membranes. Obliquely sectioned, incomplete, or poorly defined tubular profiles were excluded. All morphometric measurements were performed blinded to genotype. Measurements were collected from nonoverlapping fields. Tubular diameters between control and *Hnf4a* mutant mice were compared using two-tailed unpaired *t* tests, with convoluted and straight PTs analyzed separately. Statistical analyses were performed using GraphPad Prism version 9.0.0 (GraphPad Software, San Diego, CA).

### Kidney weight–to–body weight ratio and urinary glucose assessment

Body weight and kidney weight were recorded at two months of age. Kidney weight was normalized to body weight and expressed as a percentage using the following calculation: kidney weight/body weight × 100. Male and female mice were analyzed separately. Fresh urine samples were collected from control and *Hnf4a* mutant mice and assessed for glucose using Fisherbrand 10-SG Urine Reagent Strips (Fisher 23-111-262). Glucose levels were determined according to the manufacturer’s instructions by comparing strip color development with the reference scale provided by the manufacturer.

### Cleavage Under Targets and Release Using Nuclease (CUT&RUN)

Genome-wide mapping of HNF4A-bound sites was performed as previously described, with minor modifications.^32^ In brief, kidneys were harvested from 3-month-old mice and cells were dissociated by mincing and incubation in TrypLE Select Enzyme (Thermo A1217701) with gentle trituration. The resulting cell suspension was washed and filtered through a 40 µm cell strainer (Falcon, 352340). PT cells were immobilized on agarose-bound Lotus tetragonolobus Lectin (Vector Laboratories AL-1323), permeabilized in 0.1% digitonin, then incubated with anti-HNF4A antibody (Abcam ab41898). Recombinant Protein A-MNase (pK19p-MN, Addgene plasmid #86973) fusion protein that binds the primary antibody was added at 14 μg/ml, followed by 2mM calcium chloride addition to activate MNase enzyme, which results in cleavage and release of target DNA fragments. Library preparation and sequencing were performed as previously described for ChIP-seq.^16^

### Bioinformatic analyses

RNA-seq reads were aligned to the University of California, Santa Cruz (UCSC) mouse genome assembly mm10 using the STAR aligner.^33^ Only uniquely mapped reads were retained for downstream analysis. Raw gene-level read counts were quantified using FeatureCounts^34^ with the options “-s 2 -O —fracOverlap 0.8.” Differential gene expression analysis was conducted using DESeq2.^35^ Genes with a fold change greater than 1.5 and a false discovery rate (FDR) below 0.01 were considered differentially expressed. Gene ontology analysis was performed using EnrichR.^36^ CUT&RUN reads were aligned to mm10 using STAR,^33^ with the option “--alignSJDBoverhangMin 999 --alignIntronMax 1 --alignMatesGapMax 1000 --outFilterMultimapNmax 1 --outFilterMismatchNoverLmax 0.05 --outReadsUnmapped None --alignEndsProtrude 2 ConcordantPair". Read pairs were concatenated to form fragments. Two subsets of fragments were selected by their length: 1) nucleosome-free (NFR) < 120bp and 2) nucleosomal (NUC) > 150bp. HNF4A peak calling was performed focusing on the NFR fragments using Homer^37^ against a matching control. Peaks > 1 read per million (RPM) were retained for the downstream analysis. De novo motif search was done using Homer.^37^

## Results

### Slc34a1Cre-mediated deletion causes progressive postnatal loss of HNF4A in mature PT cells

We previously showed that *Hnf4a* is required for PT maturation, but not for initial PT specification, using Six2TGC- and Osr2Cre-mediated deletion during kidney development.^15, 16^ However, because mature PT cells fail to form in these models, they cannot determine whether HNF4A is required to maintain PT identity after differentiation. To address this question, we initially attempted inducible deletion using tamoxifen-inducible Slc34a1CreERT2.^38^ However, efficient *Hnf4a* deletion was not achieved despite repeated tamoxifen administration. We therefore used a constitutively active Slc34a1Cre allele^18^ to delete *Hnf4a* after PT maturation. Lineage tracing with the Rosa26-NuTRAP reporter demonstrated that Slc34a1Cre activity was largely restricted to HNF4A-positive, CDH6-low mature PT cells and was rarely detected in HNF4A-positive, CDH6-high immature PT cells during kidney development (Figure 1A), indicating that this model selectively targets differentiated PT cells.

**Figure 1.**
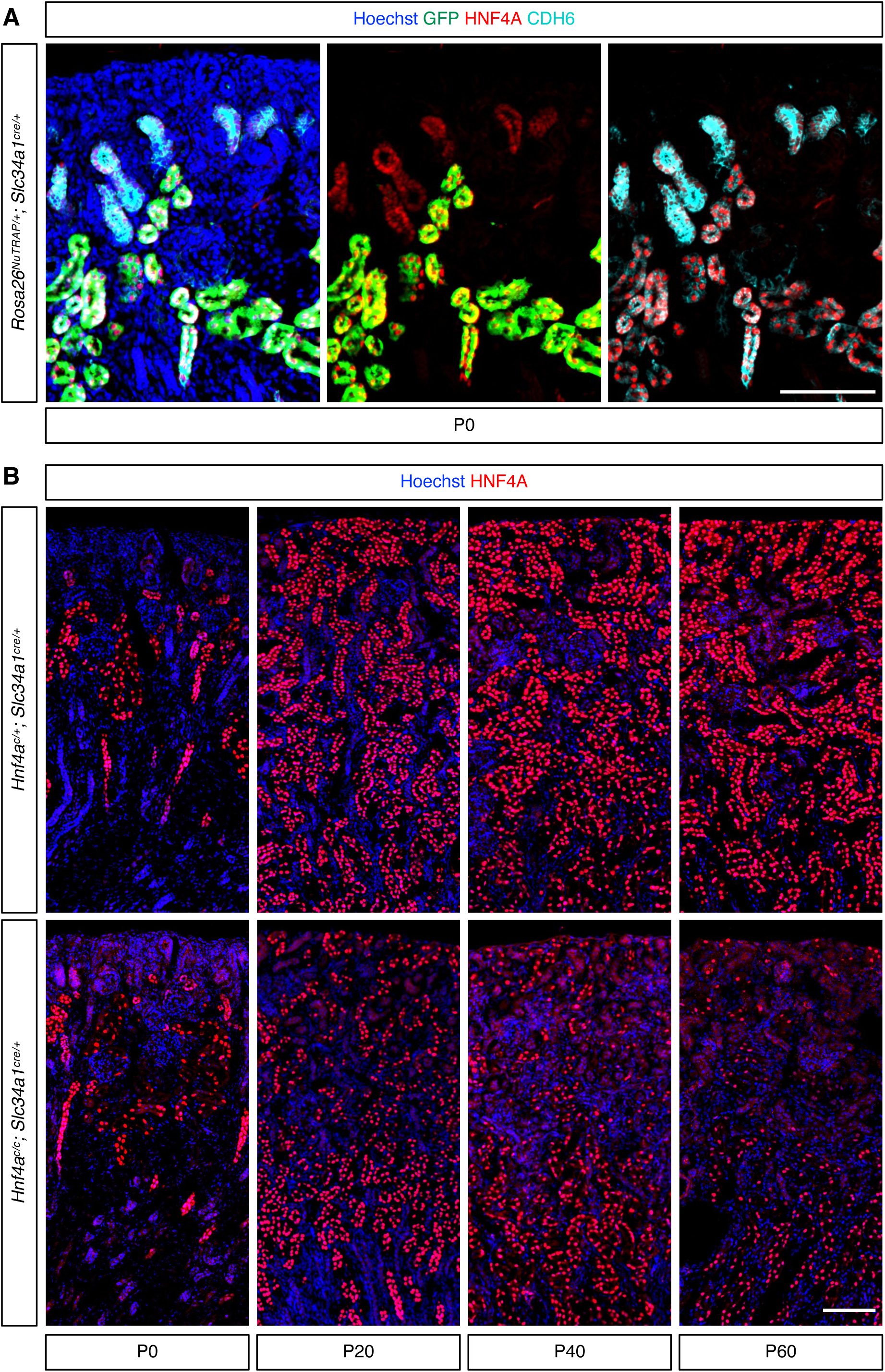
*Slc34a1Cre*-mediated deletion of *Hnf4a* causes progressive loss of HNF4A postnatally. **(A)** *Slc34a1Cre*-mediated activation of the Rosa26-NuTRAP GFP reporter is observed in mature proximal tubules (HNF4A+ CDH6-low), whereas the reporter is largely inactive in immature proximal tubules (HNF4A+ CDH6-high). Stage: P0; scale bar: 100 μm. **(B)** Loss of HNF4A following *Slc34a1Cre*-mediated deletion was undetectable at birth but became progressively evident in mutant kidneys during postnatal development. P denotes postnatal day; scale bar: 100 μm.

Although Slc34a1Cre-mediated lineage labeling was evident at postnatal day 0 (P0) (Figure 1A), HNF4A protein expression in mutant kidneys was comparable to that in controls (Figure 1B), indicating that HNF4A depletion did not occur immediately following recombination. Instead, HNF4A protein was progressively lost from PT cells during postnatal development, with depletion becoming apparent by P20 and extensive within cortical PTs by P60. In contrast, HNF4A expression was retained in straight PTs within the outer stripe of the outer medulla, likely corresponding to the S3 segment (Figure 1B). These findings demonstrate progressive postnatal depletion of HNF4A, predominantly in cortical PTs, following Slc34a1Cre-mediated deletion of *Hnf4a*.

### Loss of HNF4A occurs predominantly in convoluted proximal tubules of mutant kidneys

To confirm *Hnf4a* deletion within PTs, we examined HNF4A protein expression in lineage-labeled cells identified by the Rosa26-NuTRAP reporter activated by Slc34a1Cre. In both control and mutant kidneys, GFP-positive tubular cells were detected in convoluted and straight PT segments (Figure 2A). In control kidneys, all GFP-positive cortical PT cells retained HNF4A protein expression. In contrast, a substantial fraction of GFP-positive cortical PT cells in mutant kidneys lacked detectable HNF4A protein, confirming efficient loss of HNF4A in PT cells (Figure 2A). To assess segment-specific loss of HNF4A, we used SATB2, a marker of the S3 straight PT segment.^3^ In control kidneys, HNF4A protein was present in both SATB2-positive straight PTs and SATB2-negative convoluted PTs (Figure 2B). In mutant kidneys, SATB2-positive straight PTs largely retained HNF4A expression, whereas SATB2-negative segments, corresponding to convoluted PTs, showed marked loss of HNF4A (Figure 2B). These findings demonstrate that Slc34a1Cre-mediated deletion of *Hnf4a* occurs predominantly in convoluted PT cells.

**Figure 2.**
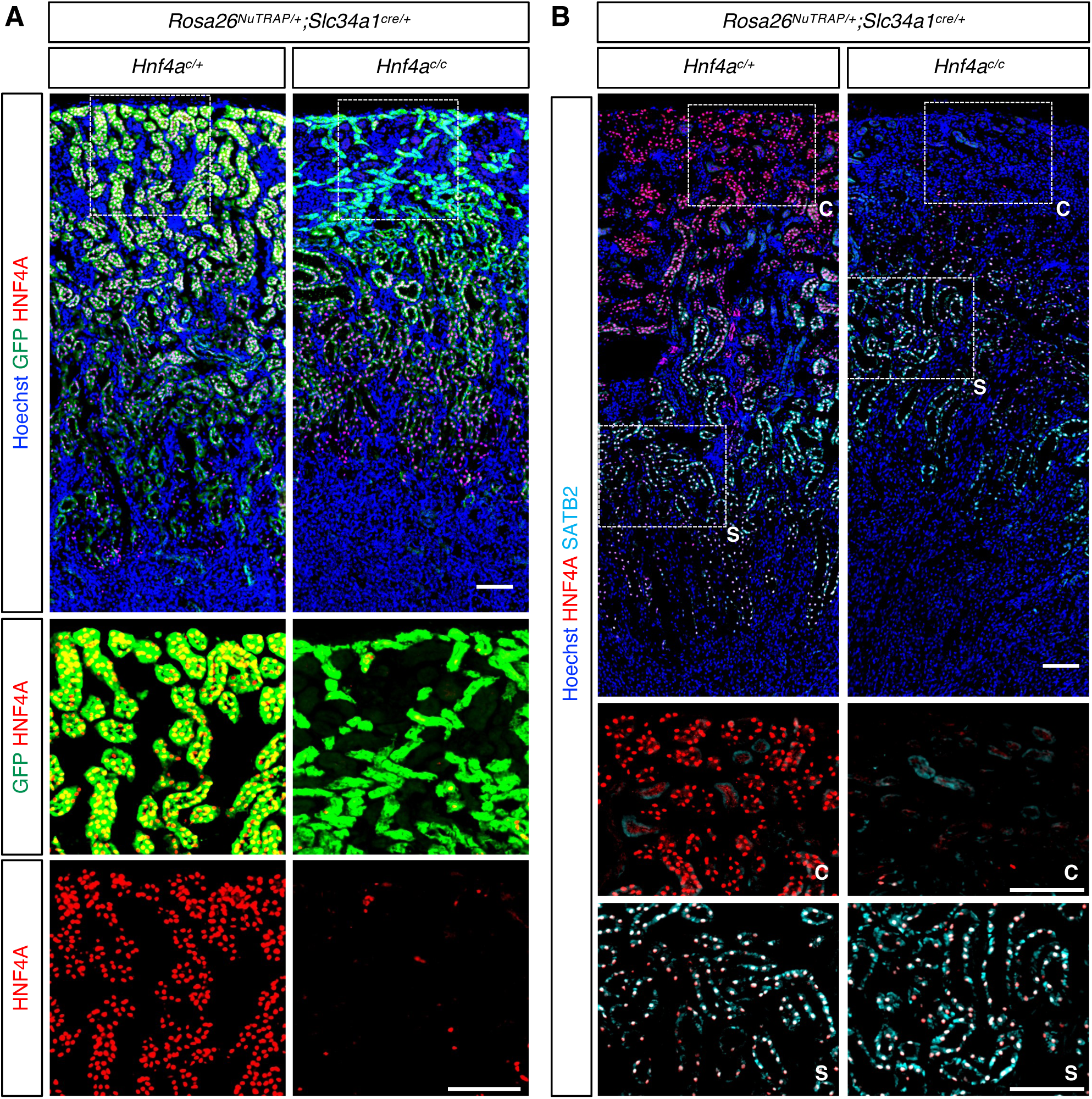
Loss of HNF4A occurs predominantly in convoluted proximal tubules of mutant kidneys. **(A)** In control and mutant kidneys, *Slc34a1Cre* activates Rosa26-NuTRAP GFP reporter expression in both convoluted and straight proximal tubules. In the cortex of control kidneys, all GFP+ cells retained HNF4A. In contrast, in the cortex of mutant kidneys, most of the GFP+ cells lack HNF4A, indicating successful deletion of *Hnf4a* in these proximal tubules. The top row shows merged Hoechst, GFP, and HNF4A staining, with the dashed box marking the region shown at higher magnification in the panels below (GFP and HNF4A merge, and HNF4A alone). Stage: P61; scale bar: 100 μm. **(B)** SATB2 is predominantly detected in straight proximal tubules. In control kidneys, HNF4A is present in both SATB2+ straight and SATB2- convoluted proximal tubules. In mutant kidneys, HNF4A is absent in SATB2- convoluted proximal tubules but largely retained in SATB2+ straight proximal tubules, indicating that HNF4A loss occurs predominantly in convoluted proximal tubules. The top row shows merged HNF4A, SATB2, and Hoechst staining, with dashed boxes marking convoluted (c) and straight (s) proximal tubule regions shown at higher magnification in the panels below. c, convoluted proximal tubule; s, straight proximal tubule. Stage: P61; scale bar: 100 μm.

### Loss of HNF4A causes selective atrophy of convoluted PTs and remodeling of renal cortical architecture

To determine the structural consequences of HNF4A loss in adult PT cells, we examined tubular morphology and epithelial organization in control and mutant kidneys. Mutant kidneys exhibited marked PT atrophy, with the most pronounced changes observed in convoluted PTs (Figure 3A). Quantification confirmed a significant reduction in the diameter of convoluted PTs in mutant kidneys, whereas the diameter of straight PTs was not significantly altered (Figure 3B). Consistent with this tubular atrophy, *Hnf4a* mutant mice exhibited significantly reduced kidney-to-body weight ratios in both males and females compared with control littermates (Supplemental Figures 1A and 1B). Mutant mice also developed glucosuria, consistent with impaired PT glucose reabsorption by convoluted PT cells (Supplemental Figure 1C). Together, these findings indicate that loss of HNF4A results in selective atrophy of convoluted PTs and impaired PT function.

**Figure 3.**
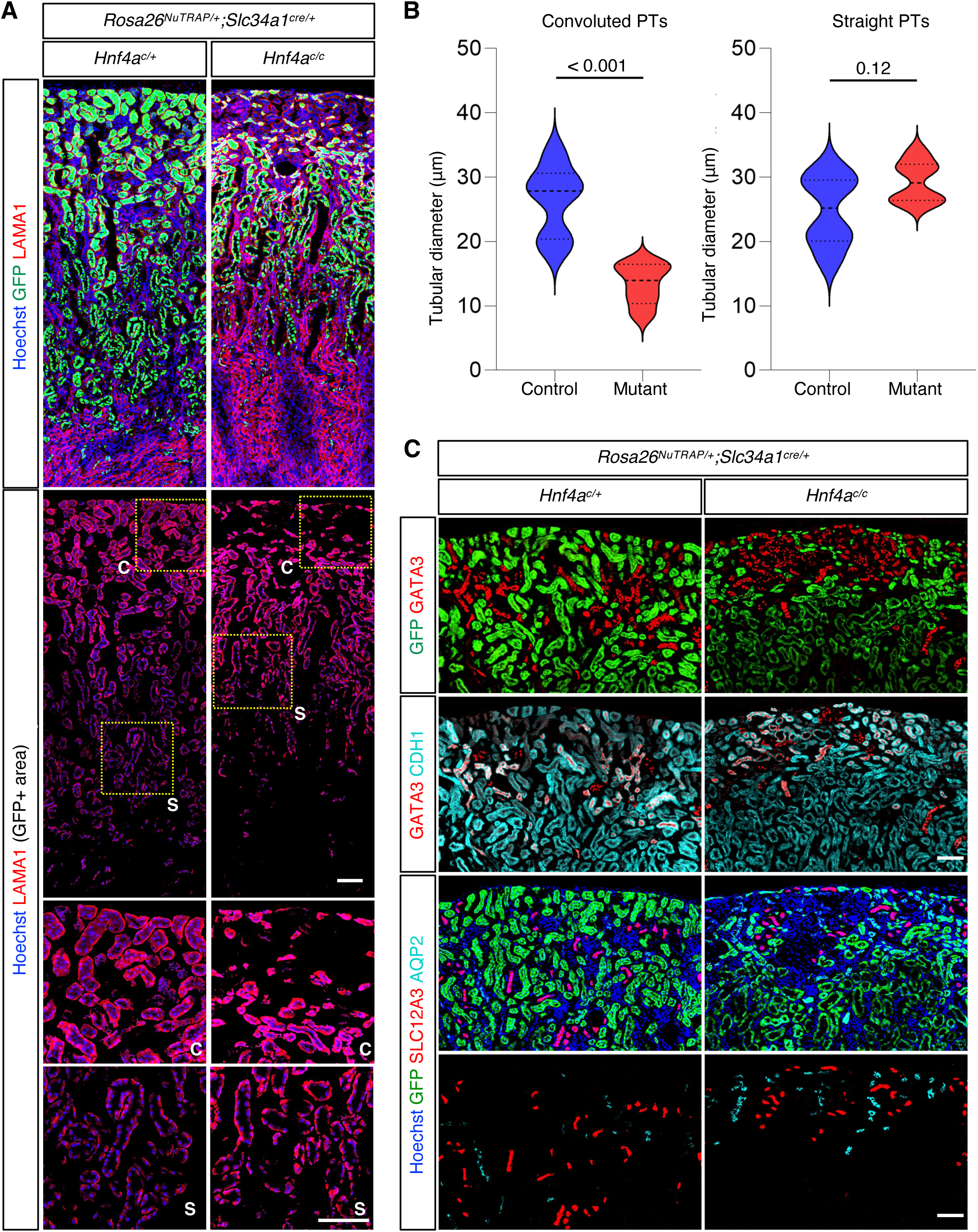
Loss of HNF4A causes selective atrophy of convoluted proximal tubules and remodeling of renal cortical architecture. **(A)** Laminin (LAMA1) staining outlines the tubular basement membrane and was used to measure tubular diameter. GFP+ (Slc34a1Cre-targeted) convoluted proximal tubules exhibit marked atrophy in Hnf4a mutant kidneys compared with controls, whereas straight proximal tubules are largely preserved. The top row shows merged GFP, LAMA1, and Hoechst staining. A GFP-based mask was then applied to restrict visualization to Slc34a1Cre-targeted proximal tubules (second row), with dashed boxes marking convoluted (c) and straight (s) proximal tubule regions shown at higher magnification in the panels below. c, convoluted proximal tubule; s, straight proximal tubule. Stage: P61; scale bar: 100 μm. **(B)** Quantification of tubular diameter in convoluted and straight proximal tubules of control and *Hnf4a* mutant kidneys. Convoluted proximal tubule diameter was lower in Hnf4a mutant kidneys than in controls (\*\*\**P* < 0.001), whereas straight proximal tubule diameter was similar between groups (ns, not significant, *P* = 0.12). Violin plots show the distribution of tubular diameters; dashed lines indicate the median and dotted lines indicate the first and third quartiles. Statistical comparisons were performed using two-tailed unpaired *t* tests (*n* = 6). **(C)** Regions exhibiting loss of GFP+ proximal tubules in *Hnf4a* mutant kidneys are associated with redistribution of GATA3+ epithelial structures toward the renal cortex (top row). Co-staining with the epithelial marker CDH1 confirms that GATA3+ cells represent distinct epithelial structures rather than non-epithelial cortical cells (second row). Co-immunostaining with SLC12A3 and AQP2 (third and fourth rows) identifies these GATA3+ structures as distal convoluted tubules (SLC12A3+) and collecting ducts (AQP2+), respectively, indicating remodeling of cortical tubular architecture following selective atrophy of convoluted proximal tubules. Stage: P61; Scale bar: 100 μm.

To determine whether PT atrophy was accompanied by changes in renal cortical organization, we examined the distribution of distal tubular markers. In mutant kidneys, cortical regions with reduced GFP-labeled PTs exhibited an increased abundance of GATA3-positive and CDH1-positive epithelial structures (Figure 3C). Co-staining with SLC12A3 and AQP2 identified these structures as distal convoluted tubules and collecting ducts, respectively. Compared with controls, distal convoluted tubules and collecting ducts occupied cortical regions normally occupied by PTs, indicating substantial remodeling of renal cortical architecture following *Hnf4a* deletion, although these data cannot distinguish whether this redistribution reflects active expansion of distal segments or passive replacement of atrophic PTs. Together, these findings demonstrate that loss of HNF4A alters the spatial organization of nephron segments within the renal cortex.

### Integrated RNA-seq and CUT&RUN analyses identify candidate direct HNF4A target genes

To define the transcriptional program directly regulated by HNF4A in adult PTs, we performed bulk RNA-seq of FACS-isolated PT cells following *Hnf4a* deletion (Supplemental Table 2) and genome-wide mapping of HNF4A-binding sites in adult mouse kidneys by CUT&RUN (Supplemental Table 3).^32, 39^ RNA-seq identified 2,176 downregulated and 2,182 upregulated genes in *Hnf4a* mutant PTs using thresholds of fold change >1.5 and FDR <0.01 (Figure 4A). Two HNF4A CUT&RUN replicates identified 10,645 reproducible binding sites. Integration of the RNA-seq and CUT&RUN datasets revealed that 1,976 (91%) of the downregulated genes and 1,524 (70%) of the upregulated genes were associated with HNF4A-binding sites (Figure 4B and Supplemental Table 4). These genes were therefore classified as candidate direct HNF4A targets.

**Figure 4.**
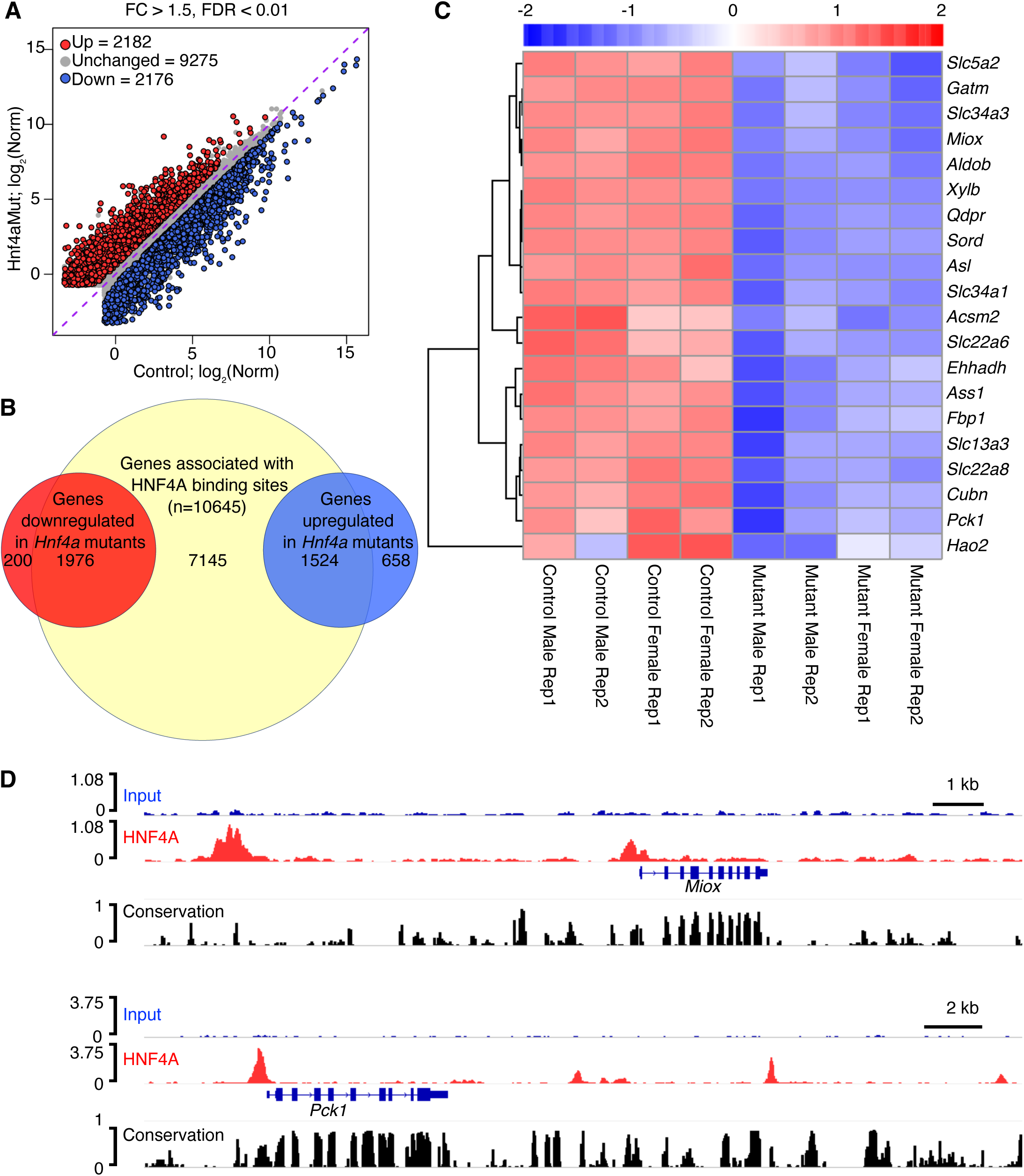
Integrated RNA-seq and CUT&RUN analyses identify direct HNF4A target genes. **(A)** Scatter plot comparing normalized gene expression levels between control and *Hnf4a* mutant kidneys determined by RNA-seq. Genes significantly upregulated (red) or downregulated (blue) in mutant kidneys relative to controls are highlighted (FC > 1.5, FDR < 0.01). The dashed line indicates equal expression between groups. Loss of *Hnf4a* results in widespread transcriptional alterations, including 2,182 upregulated and 2,176 downregulated genes. **(B)** Venn diagrams showing overlap between genes associated with HNF4A-binding sites identified by CUT&RUN and genes differentially expressed following *Hnf4a* deletion. Of the 2,176 downregulated genes, 1,976 are associated with HNF4A-binding sites, identifying them as candidate direct positive targets of HNF4A. Of the 2,182 upregulated genes, 1,524 are associated with HNF4A-binding sites. **(C)** Heatmap of representative direct HNF4A target genes identified through integration of the RNA-seq and CUT&RUN datasets. Genes involved in proximal tubule transport and metabolic functions, including *Slc5a2*, *Slc34a3*, *Miox*, *Aldob*, and *Pck1*, are downregulated in mutant kidneys relative to controls. Gene expression values are scaled by row and shown by Z-score color mapping. **(D)** Representative CUT&RUN genome browser tracks showing HNF4A occupancy at regulatory regions of *Miox* and *Pck1* genes, suggesting direct transcriptional regulation by HNF4A. Input and HNF4A CUT&RUN peaks are shown together with gene annotations and conservation tracks. Scale bars are indicated for each locus.

Among candidate direct HNF4A targets, genes involved in PT transport and metabolism were broadly downregulated following *Hnf4a* deletion, including *Slc5a2*, *Slc34a1*, *Gatm*, *Miox*, *Fbp1*, and *Pck1* (Figure 4C). CUT&RUN analysis revealed HNF4A occupancy at both promoter and enhancer regions of the *Miox* and *Pck1* loci, consistent with direct transcriptional regulation by HNF4A (Figure 4D). As additional illustrative examples, HNF4A-binding sites were identified at regulatory regions associated with *Ass1*, *Gatm*, and *Slc34a1* (Supplemental Figure 2), further supporting their classification as candidate direct HNF4A targets.

Functional enrichment analysis revealed distinct biological programs among candidate direct HNF4A targets (Supplemental Figure 3 and Supplemental Table 5). Downregulated targets were enriched for pathways related to mitochondrial function, fatty acid metabolism, solute transport, and other specialized PT functions. In contrast, upregulated HNF4A-bound genes were enriched for pathways associated with cell adhesion, extracellular matrix organization, and epithelial remodeling, consistent with activation of epithelial remodeling programs following HNF4A loss.

Motif enrichment analysis of HNF4A-bound regions revealed strong enrichment of the canonical HNF4A motif in peaks associated with both downregulated and upregulated genes (Supplemental Figure 4), confirming the specificity of the CUT&RUN dataset. Motifs recognized by several nuclear receptors, including PPAR, ESRRA, RAR, RXR, and COUP-TFII, were also enriched, consistent with shared DNA-binding preferences among nuclear receptor transcription factors. In contrast, enrichment of the structurally distinct HNF1B motif suggests potential coordinated regulation by HNF4A and HNF1B in PT cells. Together, these findings support a central role for HNF4A in maintaining the adult PT transcriptional network through direct regulation of genes required for transport and metabolic function, while limiting activation of epithelial remodeling programs.

### Loss of HNF4A induces downregulation of PT markers in convoluted PTs

To validate candidate direct HNF4A target genes identified by integrated RNA-seq and CUT&RUN analyses and to assess segment-specific alterations in PT identity, we examined their protein expression by immunofluorescence in control and mutant kidneys. In control kidneys, strong LTL staining marked convoluted PTs, whereas weaker LTL staining identified straight PTs. In *Hnf4a* mutant kidneys, LTL-high convoluted PTs were no longer detectable, whereas LTL-low straight PTs remained readily identifiable (Figure 5). Consistent with the transcriptomic data, expression of multiple PT markers was reduced in *Hnf4a* mutant kidneys, with the most pronounced changes occurring in convoluted PTs. In control kidneys, the S1-enriched markers SLC5A2 and GATM and the S2-enriched transporter SLC13A3 were detected predominantly in convoluted PTs, whereas PCK1, MIOX, ASS1, and FBP1 were expressed in both convoluted and straight PTs. In *Hnf4a* mutant kidneys, SLC5A2, GATM, and SLC13A3 staining was absent, whereas PCK1, MIOX, ASS1, and FBP1 staining was markedly reduced in convoluted PTs but largely preserved in putative straight PT segments (Figure 5) that retained HNF4A (Figure 2B). Together, these findings demonstrate that HNF4A is required to maintain differentiated PT transport and metabolic programs, and that the segment-specific pattern of marker downregulation mirrors the segment-specific loss of HNF4A itself.

**Figure 5.**
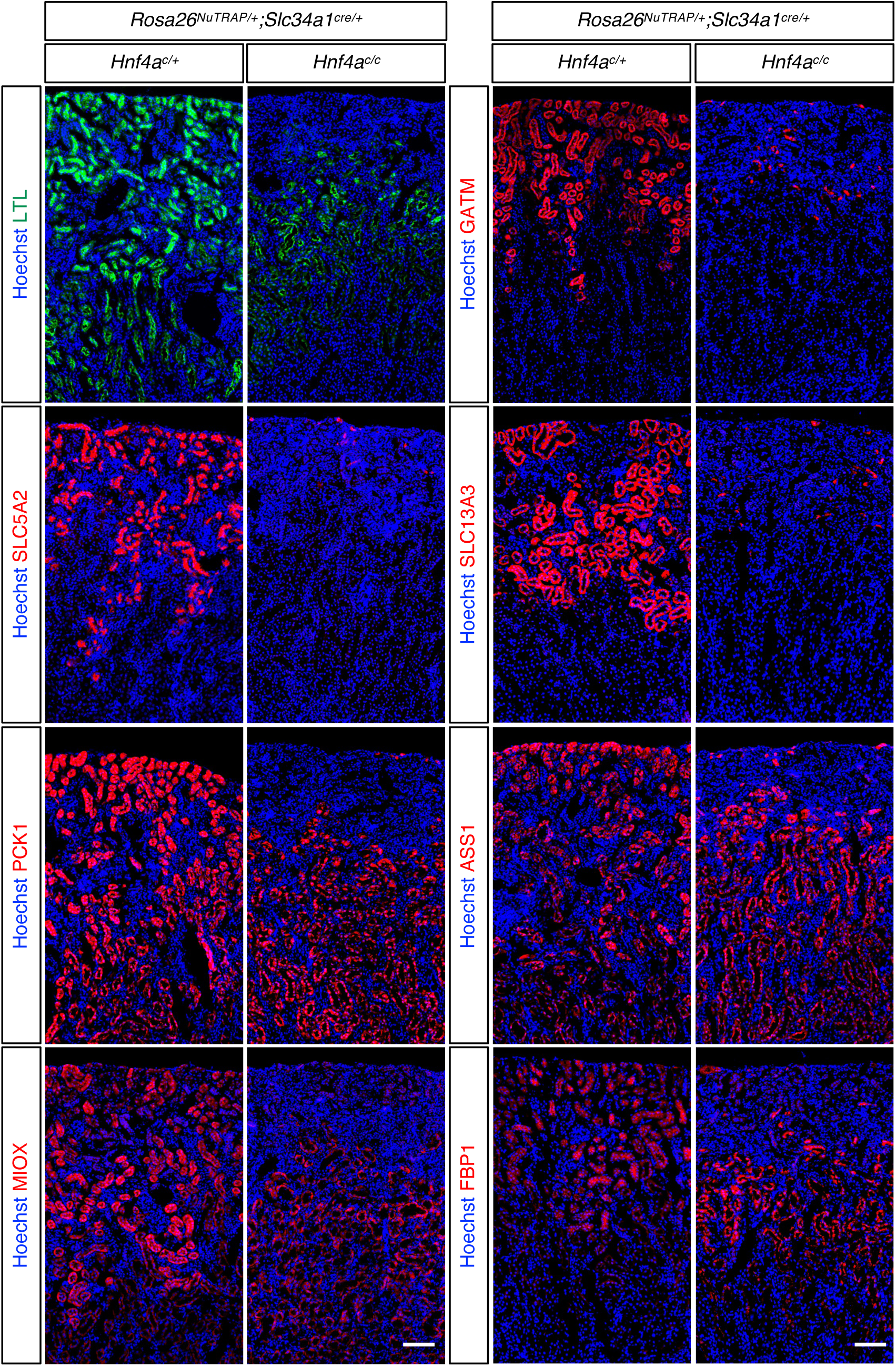
Loss of HNF4A induces segment-specific downregulation of proximal tubule markers. Representative immunofluorescence images of control and *Hnf4a* mutant kidneys stained with LTL or antibodies against SLC5A2, SLC13A3, GATM, PCK1, MIOX, ASS1, or FBP1. In control kidneys, LTL-high tubules correspond to convoluted proximal tubules, whereas LTL-low tubules represent straight proximal tubules. *Hnf4a* mutant kidneys show loss of LTL-high proximal tubules, whereas LTL-low proximal tubules remain present. SLC5A2, SLC13A3, and GATM are predominantly expressed in convoluted proximal tubules, whereas PCK1, MIOX, ASS1, and FBP1 are expressed in both convoluted and straight proximal tubules. In *Hnf4a* mutant kidneys, SLC5A2, SLC13A3, and GATM expression is lost, while PCK1, MIOX, ASS1, and FBP1 are markedly reduced in convoluted proximal tubules but largely preserved in putative straight proximal tubules. Stage: P61; scale bars: 100 μm.

### Loss of HNF4A activates injury-associated transcriptional programs in proximal tubules

RNA-seq analysis of *Hnf4a* mutant PTs revealed activation of injury-associated gene programs. To validate these findings and determine their spatial distribution, we examined the injury-associated markers VCAM1, HAVCR1, and AKAP12 by immunofluorescence. In control kidneys, VCAM1, HAVCR1, and AKAP12 were absent or detected at only minimal levels in PTs. In contrast, *Hnf4a* mutant kidneys exhibited robust induction of VCAM1 in GFP-positive PT cells that had lost HNF4A (Figure 6A). VCAM1 and HAVCR1 were both induced following *Hnf4a* deletion, although their staining patterns only partially overlapped (Figure 6B). Whereas VCAM1 expression was restricted to GFP-positive, *Hnf4a*-deficient PTs, HAVCR1 was also detected in a subset of tubules that retained HNF4A, indicating heterogeneous activation of these programs across PT segments (Figure 6C). AKAP12 was absent from control PTs but was robustly induced in GFP-positive PTs following *Hnf4a* deletion (Figure 6D).

**Figure 6.**
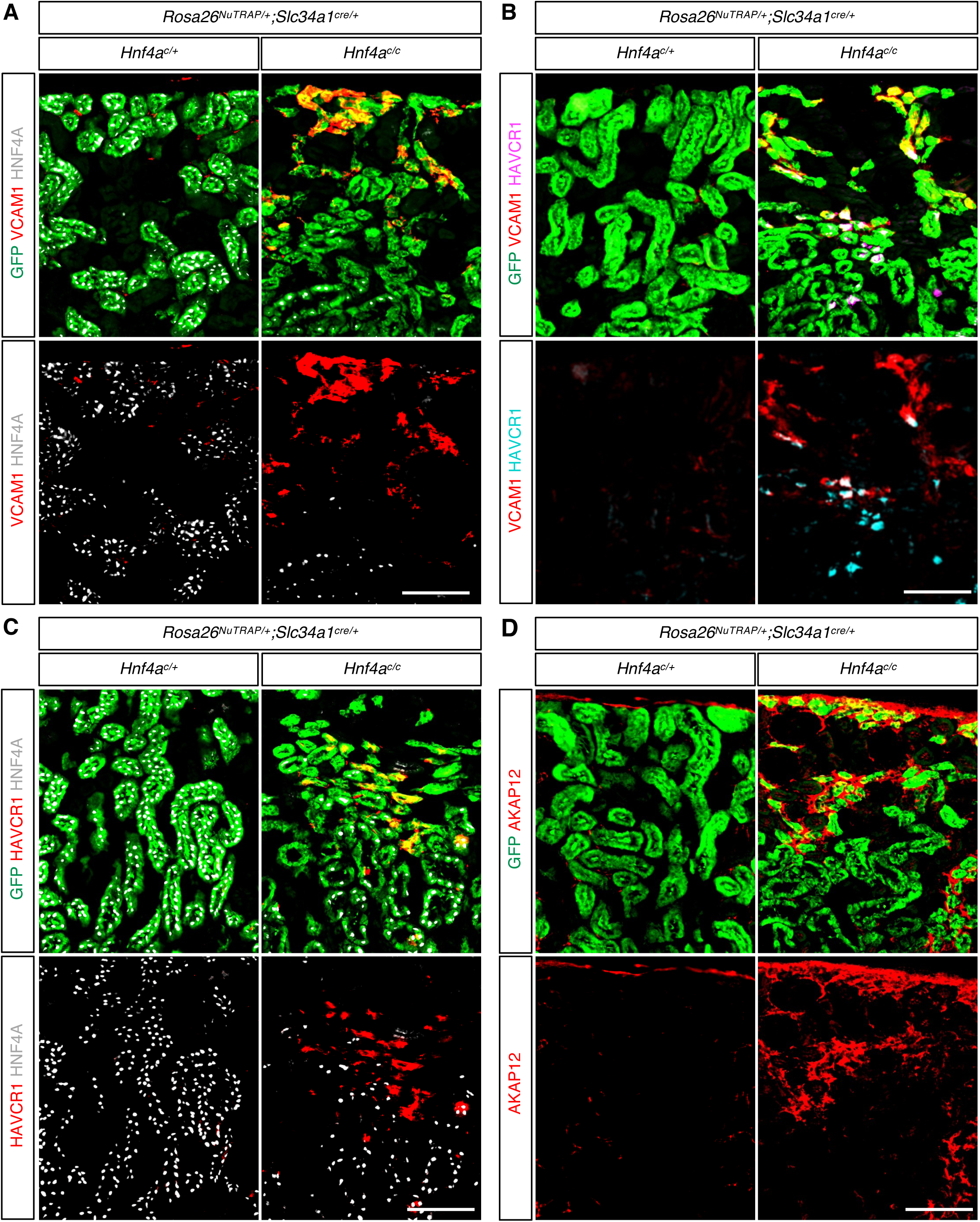
Loss of HNF4A induces injury-associated markers in proximal tubules. **(A)** GFP, HNF4A, and VCAM1 co-staining in control and *Hnf4a* mutant kidneys show that VCAM1 is not detected in control proximal tubules but is robustly induced in GFP+/HNF4A-deficient proximal tubule cells in mutant kidneys, indicating activation of an injury-associated program following HNF4A loss. Stage: P61; scale bar: 100 μm. **(B)** GFP, VCAM1, and HAVCR1 co-staining demonstrates induction of both injury markers in mutant proximal tubules. VCAM1 and HAVCR1 show partially overlapping staining patterns, indicating heterogeneous activation of injury-associated markers among proximal tubule cells. Stage: P61; scale bar: 100 μm. **(C)** GFP, HNF4A, and HAVCR1 co-staining shows induction of HAVCR1 in mutant kidneys. HAVCR1 is detected in GFP+/HNF4A-deficient proximal tubules and in a subset of neighboring tubules that retain HNF4A. Stage: P61; scale bar: 100 μm. **(D)** GFP and AKAP12 staining reveals robust AKAP12 induction in GFP+ proximal tubules following *Hnf4a* deletion, with minimal signal detected in control proximal tubules. Stage: P61; scale bar: 100 μm.

To place these findings in the context of previously defined injury-associated PT states, we compared the *Hnf4a* mutant RNA-seq dataset with injured PT genes identified by Gerhardt et al. in the Ki67-INTACT multiomic analysis of AKI repair,^40^ PT-VCAM1-related genes identified by Muto et al.,^41^ and the injured PT and FR-PTC (failed-repair proximal tubule cell) signatures described by Kirita et al. in a mouse ischemia-reperfusion injury model.^7^ *Hnf4a* mutant PTs showed significant enrichment of injured PT and PT-VCAM1 gene signatures (Supplemental Figure 5). Among the injury-associated PT states identified by Kirita et al., *Hnf4a* mutant PTs preferentially upregulated FR-PTC signature genes, whereas enrichment of the injured PT signature was not significant (Supplemental Figure 6). These findings indicate that loss of HNF4A promotes a transcriptional state that most closely resembles the FR-PTC population, which has been associated with failed repair following AKI.

Together, these findings demonstrate that loss of HNF4A activates an injury-associated transcriptional program in PT cells. This program is characterized by induction of VCAM1, HAVCR1, and AKAP12 and most closely resembles the FR-PTC state, with additional overlap with previously described injured PT and PT-VCAM1 signatures. These findings identify HNF4A as a key regulator of PT identity whose loss promotes transition toward injury-associated cell states.

### Loss of HNF4A is associated with acquisition of an immature PT-like transcriptional state

In the preceding analyses, we found that *Hnf4a*-deficient PT cells induced the injury-associated marker VCAM1. To further characterize the phenotypic state of these cells, we examined CDH6, a marker of the injury-associated FR-PTC state^7^ that we previously showed is also highly expressed in immature PT cells during kidney development but is absent from mature adult PTs under homeostatic conditions.^16^ CDH6 was co-detected with VCAM1 in GFP+ PT cells lacking HNF4A (Supplemental Figure 7A). The re-expression of CDH6 together with VCAM1 therefore raised the possibility that *Hnf4a*-deficient PT cells not only activate injury-associated programs but also acquire features of an immature PT state.

We next examined LRP2, which remained detectable in a subset of GFP+ *Hnf4a*-deficient PT cells despite the broad reduction in other mature PT markers following *Hnf4a* deletion (Supplemental Figure 7B). To interpret this finding, we analyzed our E18.5/P0 reference scRNA-seq dataset containing immature and mature PT populations (*Supplemental Figure 7C*).^18, 42^ Feature-plots and violin-plots showed that the mature PT markers *Gatm*, *Miox*, and *Pck1* were expressed predominantly in mature PTs, whereas *Cdh6* and *Vcam1* were enriched in immature PTs (Supplemental Figure 7D). In contrast, *Lrp2* and *Hnf4a* were detected in both immature and mature PT populations, although their expression was higher in mature PTs. These findings support the association of *Cdh6* and *Vcam1* with immature PT states while demonstrating that *Lrp2* is expressed across both immature and mature PT populations.

To determine whether the co-detection of CDH6 and VCAM1 reflected a developmental expression pattern, we examined their distribution during kidney development. Immature PT cells, identified by HNF4A+ cells with strong CDH6 signal, differentiate into mature PT cells, which are characterized by HNF4A+ cells with weak CDH6 signal (Supplemental Figure 8).^16, 43^ VCAM1 was detected in a subset of immature PT cells in the developing kidney, whereas it was absent from mature PTs (Supplemental Figure 8). These observations support the conclusion that *Hnf4a*-deficient adult PTs acquire molecular features normally associated with immature PT cells.

To define this transcriptional shift more broadly, we first filtered the immature and mature PT marker lists from the E18.5/P0 reference scRNA-seq dataset^18, 42^ using thresholds of P < 0.01 and log_2_FC > 1.5 relative to other cell populations. We then retained genes that were either present only in the immature PT marker list or, when present in both marker lists, exhibited higher average expression in immature PTs than in mature PTs. This analysis identified 52 immature PT-associated genes. Of these, 48 were detected in the bulk RNA-seq dataset comparing control and *Hnf4a* mutant PTs, including 27 that were upregulated, 5 that were downregulated, and 16 that were not significantly changed; the remaining 4 genes were not detected (Supplemental Table 6). Heatmap visualization of the 32 differentially expressed genes showed predominant induction of immature PT-associated genes in *Hnf4a* mutant PTs (Supplemental Figure 9). Upregulated genes included *Cdh6*, *Cryab*, *Vcam1*, *Grid1*, *Sema3c*, *Adamts1*, *Adamts16*, *Myo5b*, *Slc39a8*, *Plau*, *Dmd*, and *Phgdh*. Several of the induced genes, including *Cryab*, *Vcam1*, *Adamts1*, *Sema3c*, *Grid1*, *Myo5b*, and *Cdh6*, were also represented in one or more of the injury-associated PT gene signatures analyzed above (Supplemental Figures 5 and 6). Together, these findings indicate that loss of HNF4A is associated not only with suppression of mature PT transport and metabolic programs but also with acquisition of an immature PT-like transcriptional state that partially overlaps with the injury-associated response.

## Discussion

In this study, we demonstrate that HNF4A is required for maintenance of PT identity and function in the postnatal kidney. Although previous studies established that HNF4A is essential for PT maturation during development,^15, 16^ whether it is required to maintain the differentiated PT state remained unknown. Using Slc34a1Cre-mediated deletion of *Hnf4a* after PT maturation, we show that loss of HNF4A leads to downregulation of PT-specific transport and metabolic genes, accompanied by tubular atrophy, cortical remodeling, and activation of injury-associated transcriptional states. *Hnf4a*-deficient PT cells also acquired molecular features associated with immature and failed-repair PT populations. Together, these findings establish HNF4A as a critical regulator of adult PT homeostasis and demonstrate that maintenance of PT identity is an active process requiring continuous transcriptional regulation after differentiation.

Our findings extend previous studies defining the developmental role of HNF4A in PT differentiation. Earlier deletion of *Hnf4a* using Six2TGC or Osr2Cre prevented formation of mature PT cells, demonstrating that HNF4A is required for PT maturation.^15, 16^ As a result, it remained unclear whether the loss of PT-specific gene expression in those *Hnf4a* mutant kidneys reflected failure of differentiation or loss of HNF4A-dependent transcriptional programs. By deleting *Hnf4a* after PT maturation, the present study separates these functions and demonstrates that mature PT identity remains dependent on continued HNF4A activity. Thus, HNF4A functions not only as a developmental regulator of PT maturation but also as a PT identity-maintaining factor required to preserve the differentiated state of adult PT cells (Figures 1 and 4).

An unexpected feature of this model was the delayed loss of HNF4A, which occurred predominantly in convoluted PTs, following Slc34a1Cre-mediated deletion. Although Cre activity was evident at birth (Figure 1A), HNF4A protein persisted initially and declined progressively during postnatal life, with loss concentrated in cortical convoluted PTs while many straight PTs retained detectable HNF4A protein (Figures 1 and 2). The basis for the delayed loss of HNF4A remains unclear but may reflect differences in recombination efficiency between the *Hnf4a* and reporter loci or slow turnover of HNF4A protein. The segment-specific pattern may instead reflect regional variation in Slc34a1Cre activity; because *Slc34a1* expression is restricted to convoluted PTs in the adult kidney,^3^ Slc34a1Cre-mediated recombination at the *Hnf4a* locus may be more efficient in this segment, which could account for the preferential loss of HNF4A observed there. Mutant kidneys exhibited selective atrophy of convoluted PTs, whereas straight PTs were relatively preserved (Figures 2 and 3). PT atrophy was accompanied by redistribution of distal convoluted tubules and collecting ducts into cortical regions normally occupied by PTs, as well as development of glucosuria and reduced kidney mass (Figure 3 and Supplemental Figure 1). Although the present analyses do not distinguish active expansion of distal nephron segments from passive redistribution following PT atrophy, these findings demonstrate that disruption of HNF4A-dependent programs has substantial structural and functional consequences for the kidney.

Integration of RNA-seq and CUT&RUN datasets identified a broad network of candidate direct HNF4A target genes underlying mature PT function. A substantially greater proportion of downregulated genes than upregulated genes were associated with HNF4A binding, supporting a predominant role for HNF4A as a transcriptional activator in adult PT cells (Figure 4A,B). Direct HNF4A targets were enriched for genes involved in solute transport, mitochondrial metabolism, fatty acid utilization, gluconeogenesis, and amino acid metabolism (Figure 4C,D; Supplemental Figures 2 and 3). Consistent with these findings, loss of HNF4A resulted in coordinated suppression of transporters and metabolic enzymes at both the transcript and protein levels (Figures 4 and 5). Impaired fatty acid oxidation in renal tubular epithelial cells has been linked to tubular dedifferentiation and kidney fibrosis.^44^ Motif analysis further identified enrichment of canonical HNF4A-binding sequences and additional nuclear receptor motifs, suggesting that HNF4A functions within a broader regulatory network controlling PT differentiation and metabolism (Supplemental Figure 4). Together, these observations identify HNF4A as a central node within the transcriptional network that sustains mature PT transport and metabolic function.

A major finding of this study is that loss of HNF4A is sufficient to induce a substantial component of the injury-associated transcriptional program in the absence of exogenous injury. *Hnf4a*-deficient PT cells exhibited increased expression of *Vcam1*, *Havcr1*, *Akap12*, and other markers associated with injured and dedifferentiated PT cells (Figure 6). In addition, mutant PTs showed significant enrichment of published injury-associated signatures, including the PT-VCAM1 program and the FR-PTC state (Supplemental Figures 5 and 6). The similarity to FR-PTC is particularly notable because this population has been linked to persistent injury and incomplete restoration of differentiated PT function following AKI.^7, 8, 40, 45, 46^ Importantly, maladaptive PT states characterized by dedifferentiation and pro-inflammatory and profibrotic programs have also been identified in human AKI across diverse etiologies.^47^

*Hnf4a*-deficient PTs also reactivated genes normally associated with immature PT states, including *Cdh6*, while retaining selected features of differentiated PTs (Supplemental Figures 7-9). These findings indicate that *Hnf4a*-deficient cells do not simply revert to an early developmental state but instead acquire a hybrid phenotype characterized by loss of mature transport and metabolic functions, persistence of select PT features, and activation of both developmental and injury-associated programs. At the same time, HNF4A loss did not fully recapitulate the injury response. Notably, *Sox9*, a key injury-responsive transcription factor that is robustly induced after AKI,^48–50^ was not activated in *Hnf4a*-deficient PTs (Supplemental Table 2). This observation suggests that suppression of HNF4A is sufficient to reproduce many features of PT dedifferentiation and failed repair but is not sufficient to generate the complete injury-response program. Rather, injury-associated remodeling likely requires cooperation between loss of HNF4A-dependent differentiation programs and additional signals generated by tissue injury, inflammation, or cellular stress.

Collectively, our findings identify HNF4A as a central regulator of mature PT identity that sustains differentiated transport and metabolic programs while limiting activation of immature and injury-associated states. The observation that HNF4A loss alone is sufficient to induce many molecular features of failed-repair PT cells suggests that suppression of HNF4A may contribute directly to epithelial dysfunction during AKI and CKD progression. Although the present model does not establish whether *Hnf4a* downregulation is an initiating event or a consequence of kidney injury, it demonstrates that loss of HNF4A activity is sufficient to drive substantial components of the maladaptive PT transcriptional program. Future studies should determine whether preservation or restoration of HNF4A-dependent pathways promotes adaptive tubular repair and limits progression to chronic kidney disease.

## Acknowledgments

We thank the Robert H. Lurie Comprehensive Cancer Center of Northwestern University in Chicago, IL, for the use of the Flow Cytometry Core Facility, which provided cell sorting services. The Lurie Cancer Center is supported in part by NCI Cancer Center Support Grant P30 CA060553. We also thank the Northwestern University NUSeq Core Facility for sequencing services.

## Author Contributions

Conceptualization: Fariba Nosrati, Joo-Seop Park

Data curation: Fariba Nosrati, Joo-Seop Park

Formal analysis: Fariba Nosrati, Christopher Ahn, Hee-Woong Lim

Funding acquisition: Joo-Seop Park

Investigation: Fariba Nosrati, Zeinab Dehghani-Ghobadi, Eunah Chung, Joo-Seop Park

Methodology: Fariba Nosrati, Zeinab Dehghani-Ghobadi, Eunah Chung, Joo-Seop Park

Project administration: Joo-Seop Park

Resources: Joo-Seop Park

Software: Christopher Ahn, Hee-Woong Lim

Validation: Fariba Nosrati, Joo-Seop Park

Visualization: Fariba Nosrati, Joo-Seop Park

Writing – original draft: Fariba Nosrati, Joo-Seop Park

Writing – review & editing: Fariba Nosrati, Eunah Chung, Joo-Seop Park

## Data Sharing Statement

All transcriptomic and genomic datasets generated in this study have been deposited in the NCBI Gene Expression Omnibus (GEO). Raw data from the bulk RNA-seq and CUT&RUN experiments are available under accession numbers GSE336439 and GSE336441, respectively.

## Declaration of generative AI and AI-assisted technologies in the manuscript preparation process

During the preparation of this work, the author(s) used Copilot and Claude to assist with manuscript editing, and refinement of scientific writing. The author(s) reviewed and edited the output as needed and take full responsibility for the content of the published article.

## References

1. Wessely O, Cerqueira DM, Tran U, et al. The bigger the better: determining nephron size in kidney. Pediatr Nephrol 2014; 29: 525–530.

2. Lee JW, Chou CL, Knepper MA. Deep Sequencing in Microdissected Renal Tubules Identifies Nephron Segment-Specific Transcriptomes. J Am Soc Nephrol 2015.

3. Chen L, Chou CL, Knepper MA. A Comprehensive Map of mRNAs and Their Isoforms across All 14 Renal Tubule Segments of Mouse. J Am Soc Nephrol 2021; 32: 897–912.

4. Chevalier RL. The proximal tubule is the primary target of injury and progression of kidney disease: role of the glomerulotubular junction. Am J Physiol Renal Physiol 2016; 311: F145–161.

5. McMahon AP. Development of the Mammalian Kidney. Curr Top Dev Biol 2016; 117: 31–64.

6. Chiba T, Peasley KD, Cargill KR, et al. Sirtuin 5 Regulates Proximal Tubule Fatty Acid Oxidation to Protect against AKI. J Am Soc Nephrol 2019; 30: 2384–2398.

7. Kirita Y, Wu H, Uchimura K, et al. Cell profiling of mouse acute kidney injury reveals conserved cellular responses to injury. Proc Natl Acad Sci U S A 2020; 117: 15874–15883.

8. Gerhardt LMS, Liu J, Koppitch K, et al. Single-nuclear transcriptomics reveals diversity of proximal tubule cell states in a dynamic response to acute kidney injury. Proceedings of the National Academy of Sciences 2021; 118.

9. Lake BB, Menon R, Winfree S, et al. An atlas of healthy and injured cell states and niches in the human kidney. Nature 2023; 619: 585–594.

10. Bonventre JV, Yang L. Cellular pathophysiology of ischemic acute kidney injury. J Clin Invest 2011; 121: 4210–4221.

11. Yang L, Besschetnova TY, Brooks CR, et al. Epithelial cell cycle arrest in G2/M mediates kidney fibrosis after injury. Nat Med 2010; 16: 535–543, 531p following 143.

12. Ferenbach DA, Bonventre JV. Mechanisms of maladaptive repair after AKI leading to accelerated kidney ageing and CKD. Nat Rev Nephrol 2015; 11: 264–276.

13. Takaori K, Nakamura J, Yamamoto S, et al. Severity and Frequency of Proximal Tubule Injury Determines Renal Prognosis. J Am Soc Nephrol 2016; 27: 2393–2406.

14. Chen L, Luo S, Dupre A, et al. The nuclear receptor HNF4 drives a brush border gene program conserved across murine intestine, kidney, and embryonic yolk sac. Nat Commun 2021; 12: 2886.

15. Marable SS, Chung E, Adam M, et al. Hnf4a deletion in the mouse kidney phenocopies Fanconi renotubular syndrome. JCI Insight 2018; 3.

16. Marable SS, Chung E, Park JS. Hnf4a is required for the development of Cdh6-expressing progenitors into proximal tubules in the mouse kidney. J Am Soc Nephrol 2020.

17. Yoshimura Y, Muto Y, Omachi K, et al. Elucidating the Proximal Tubule HNF4A Gene Regulatory Network in Human Kidney Organoids. J Am Soc Nephrol 2023; 34: 1672–1686.

18. Chung E, Nosrati F, Adam M, et al. Proximal Tubule Cells Contribute to the Thin Descending Limb of the Loop of Henle during Mouse Kidney Development. J Am Soc Nephrol 2025.

19. Park JS, Valerius MT, McMahon AP. Wnt/beta-catenin signaling regulates nephron induction during mouse kidney development. Development 2007; 134: 2533–2539.

20. Kobayashi A, Valerius MT, Mugford JW, et al. Six2 defines and regulates a multipotent self-renewing nephron progenitor population throughout mammalian kidney development. Cell Stem Cell 2008; 3: 169–181.

21. Lan Y, Wang Q, Ovitt CE, et al. A unique mouse strain expressing Cre recombinase for tissue-specific analysis of gene function in palate and kidney development. Genesis 2007; 45: 618–624.

22. Clark AJ, Saade MC, Vemireddy V, et al. Hepatocyte nuclear factor 4alpha mediated quinolinate phosphoribosylltransferase (QPRT) expression in the kidney facilitates resilience against acute kidney injury. Kidney Int 2023; 104: 1150–1163.

23. Chang-Panesso M, Kadyrov FF, Lalli M, et al. FOXM1 drives proximal tubule proliferation during repair from acute ischemic kidney injury. J Clin Invest 2019; 129: 5501–5517.

24. Balzer MS, Doke T, Yang YW, et al. Single-cell analysis highlights differences in druggable pathways underlying adaptive or fibrotic kidney regeneration. Nat Commun 2022; 13: 4018.

25. Kha M, Magnusson Y, Johansson I, et al. Injured Proximal Tubular Epithelial Cells Lose Hepatocyte Nuclear Factor 4alpha Expression Crucial for Brush Border Formation and Transport. Am J Pathol 2025; 195: 845–860.

26. Rudman-Melnick V, Adam M, Potter A, et al. Single-Cell Profiling of AKI in a Murine Model Reveals Novel Transcriptional Signatures, Profibrotic Phenotype, and Epithelial-to-Stromal Crosstalk. J Am Soc Nephrol 2020; 31: 2793–2814.

27. Telang AC, Ference-Salo JT, McElliott MC, et al. Sustained alterations in proximal tubule gene expression in primary culture associate with HNF4A loss. Sci Rep 2024; 14: 22927.

28. Jiang G, Lu X, Cao R, et al. HNF4A P2 isoform alleviates kidney fibrosis by inhibiting dedifferentiation of proximal tubular cells through JAG1/NOTCH signaling. Cell Mol Biol Lett 2026; 31.

29. Roh HC, Tsai LT, Lyubetskaya A, et al. Simultaneous Transcriptional and Epigenomic Profiling from Specific Cell Types within Heterogeneous Tissues In Vivo. Cell reports 2017; 18: 1048–1061.

30. Hayhurst GP, Lee YH, Lambert G, et al. Hepatocyte nuclear factor 4alpha (nuclear receptor 2A1) is essential for maintenance of hepatic gene expression and lipid homeostasis. Mol Cell Biol 2001; 21: 1393–1403.

31. Dehghani-Ghobadi Z, Chung E, Sayed M, et al. Constitutive YAP activation in distal nephron segments disrupts epithelial identity and nephron patterning. JCI Insight 2026.

32. Skene PJ, Henikoff JG, Henikoff S. Targeted in situ genome-wide profiling with high efficiency for low cell numbers. Nat Protoc 2018; 13: 1006–1019.

33. Dobin A, Davis CA, Schlesinger F, et al. STAR: ultrafast universal RNA-seq aligner. Bioinformatics 2013; 29: 15–21.

34. Liao Y, Smyth GK, Shi W. featureCounts: an efficient general purpose program for assigning sequence reads to genomic features. Bioinformatics 2014; 30: 923–930.

35. Love MI, Huber W, Anders S. Moderated estimation of fold change and dispersion for RNA-seq data with DESeq2. Genome Biol 2014; 15: 550.

36. Kuleshov MV, Jones MR, Rouillard AD, et al. Enrichr: a comprehensive gene set enrichment analysis web server 2016 update. Nucleic Acids Res 2016; 44: W90–97.

37. Heinz S, Benner C, Spann N, et al. Simple combinations of lineage-determining transcription factors prime cis-regulatory elements required for macrophage and B cell identities. Mol Cell 2010; 38: 576–589.

38. Kusaba T, Lalli M, Kramann R, et al. Differentiated kidney epithelial cells repair injured proximal tubule. Proc Natl Acad Sci U S A 2014; 111: 1527–1532.

39. Skene PJ, Henikoff S. An efficient targeted nuclease strategy for high-resolution mapping of DNA binding sites. eLife 2017; 6.

40. Gerhardt LMS, Koppitch K, van Gestel J, et al. Lineage Tracing and Single-Nucleus Multiomics Reveal Novel Features of Adaptive and Maladaptive Repair after Acute Kidney Injury. J Am Soc Nephrol 2023; 34: 554–571.

41. Muto Y, Wilson PC, Ledru N, et al. Single cell transcriptional and chromatin accessibility profiling redefine cellular heterogeneity in the adult human kidney. Nat Commun 2021; 12: 2190.

42. Rudman-Melnick V, Adam M, Stowers K, et al. Single-cell sequencing dissects the transcriptional identity of activated fibroblasts and identifies novel persistent distal tubular injury patterns in kidney fibrosis. Sci Rep 2024; 14: 439.

43. Deacon P, Concodora CW, Chung E, et al. β-catenin regulates the formation of multiple nephron segments in the mouse kidney. Scientific Reports 2019; 9: 15915.

44. Kang HM, Ahn SH, Choi P, et al. Defective fatty acid oxidation in renal tubular epithelial cells has a key role in kidney fibrosis development. Nat Med 2015; 21: 37–46.

45. Ledru N, Wilson PC, Muto Y, et al. Predicting proximal tubule failed repair drivers through regularized regression analysis of single cell multiomic sequencing. Nat Commun 2024; 15: 1291.

46. Melchinger I, Guo K, Li X, et al. VCAM-1 mediates proximal tubule-immune cell cross talk in failed tubule recovery during AKI-to-CKD transition. Am J Physiol Renal Physiol 2024; 327: F610–F622.

47. Wen Y, Su E, Xu L, et al. Analysis of the human kidney transcriptome and plasma proteome identifies markers of proximal tubule maladaptation to injury. Sci Transl Med 2023; 15: eade7287.

48. Aggarwal S, Wang Z, Rincon Fernandez Pacheco D, et al. SOX9 switch links regeneration to fibrosis at the single-cell level in mammalian kidneys. Science 2024; 383: eadd6371.

49. Kang HM, Huang S, Reidy K, et al. Sox9-Positive Progenitor Cells Play a Key Role in Renal Tubule Epithelial Regeneration in Mice. Cell reports 2016; 14: 861–871.

50. Kumar S, Liu J, Pang P, et al. Sox9 Activation Highlights a Cellular Pathway of Renal Repair in the Acutely Injured Mammalian Kidney. Cell reports 2015.

