## Supplementary material for "HNF4A maintains proximal tubule identity and limits injury-associated cell states in the adult mouse kidney": Supp Figures

**TABLE OF CONTENTS OF SUPPLEMENTAL MATERIAL**

Supplemental Figure 1. Loss of HNF4A in convoluted proximal tubules reduces kidney mass and results in glucosuria.

Supplemental Figure 2. Representative CUT&RUN genome browser tracks show HNF4A binding at direct target gene loci.

Supplemental Figure 3. Gene Ontology and KEGG pathway enrichment analyses of candidate direct HNF4A target genes identified by integrated RNA-seq and CUT&RUN.

Supplemental Figure 4. HNF4A CUT&RUN peaks associated with differentially expressed genes are enriched for HNF4A and other transcription factor motifs.

Supplemental Figure 5. Hnf4a mutant proximal tubules show enrichment of the Injured PT and PT-VCAM1 gene signatures.

Supplemental Figure 6. Hnf4a mutant proximal tubules show preferential upregulation of the FR-PTC gene signature compared with the injured PT signature.

Supplemental Figure 7. Loss of HNF4A is associated with acquisition of an immature proximal tubule–like state.

Supplemental Figure 8. VCAM1 is detected in a subset of CDH6-high proximal tubule progenitor cells in the developing kidney.

Supplemental Figure 9. HNF4A-deficient proximal tubules acquire an immature proximal tubule–like transcriptional state.

Supplemental Table 1. Primary and secondary antibodies and lectin reagent used in this study.

Supplemental Table 2. Differential gene expression in control and *Hnf4a* mutant proximal tubules (RNA-seq).

Supplemental Table 3. HNF4A CUT&RUN analysis.

Supplemental Table 4. Genes associated with HNF4A CUT&RUN peaks and their differential expression following Hnf4a deletion in proximal tubules.

Supplemental Table 5. GO and KEGG pathway enrichment analyses of candidate direct HNF4A target genes.

Supplemental Table 6. Expression of immature PT-associated genes in *Hnf4a* mutant proximal tubules.

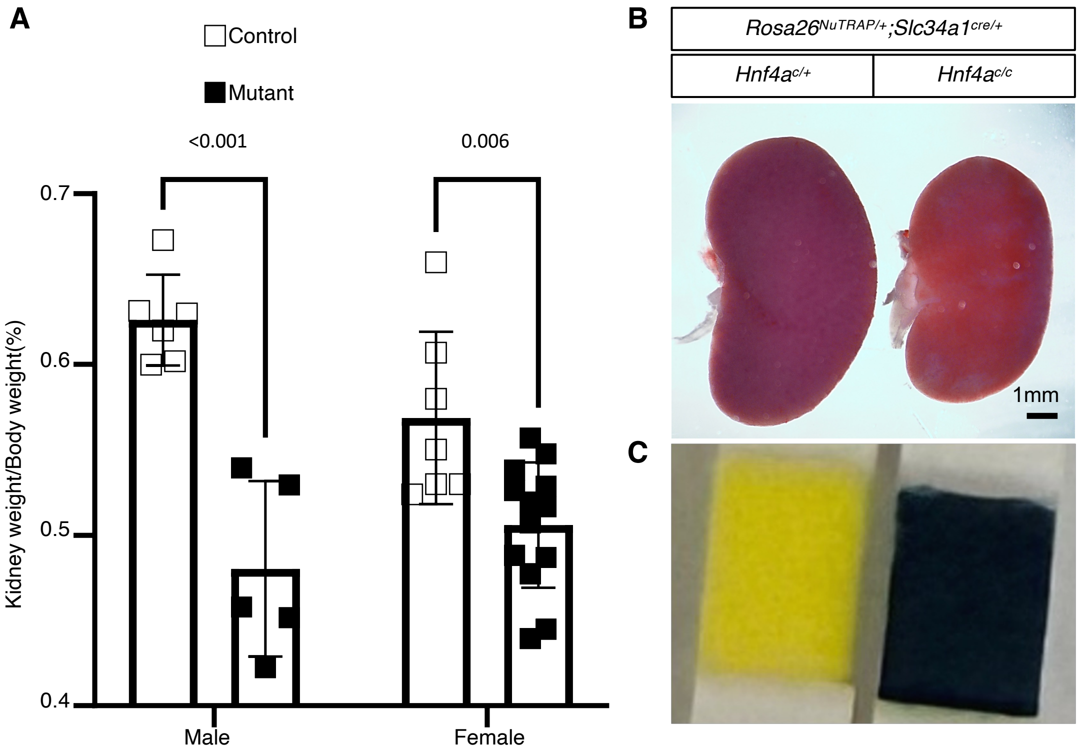


**Supplemental Figure 1. Loss of HNF4A in convoluted proximal tubules reduces kidney mass and results in glucosuria.** (A) Kidney-to-body weight ratios in male and female control and *Hnf4a* mutant mice. Mutant mice showed significantly reduced kidney-to-body weight ratios in both sexes. Individual animals are shown as open squares (control) or filled squares (mutant), with bars representing mean ± SD. Male: control, $n$ = 6; mutant, $n$ = 5. Female: control, $n$ = 7; mutant, $n$ = 13. $P < 0.001$ for males and $P = 0.006$ for females. (B) Representative kidneys from control and *Hnf4a* mutant mice showing reduced kidney size in mutants. Stage: P61; scale bar: 1 mm. (C) Representative urine glucose test strips showing glucosuria in *Hnf4a* mutant mice. Stage: P61.

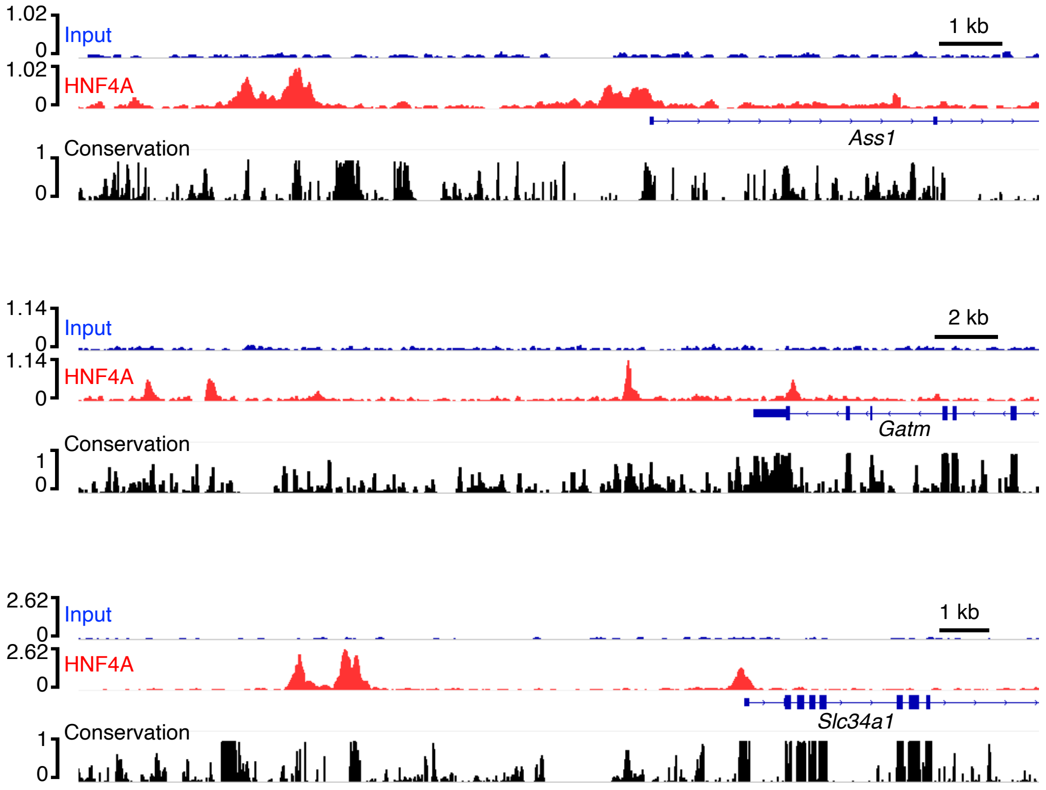


**Supplemental Figure 2. Representative CUT&RUN genome browser tracks show HNF4A binding at direct target gene loci.** Representative CUT&RUN genome browser tracks demonstrating HNF4A occupancy at regulatory regions of *Ass1*, *Gatm*, and *Slc34a1*. Input tracks are shown above corresponding HNF4A CUT&RUN tracks, and corresponding chromatin accessibility profiles are displayed below. These data further support direct binding of HNF4A at representative proximal tubule target gene loci. Scale bars are indicated for each locus.

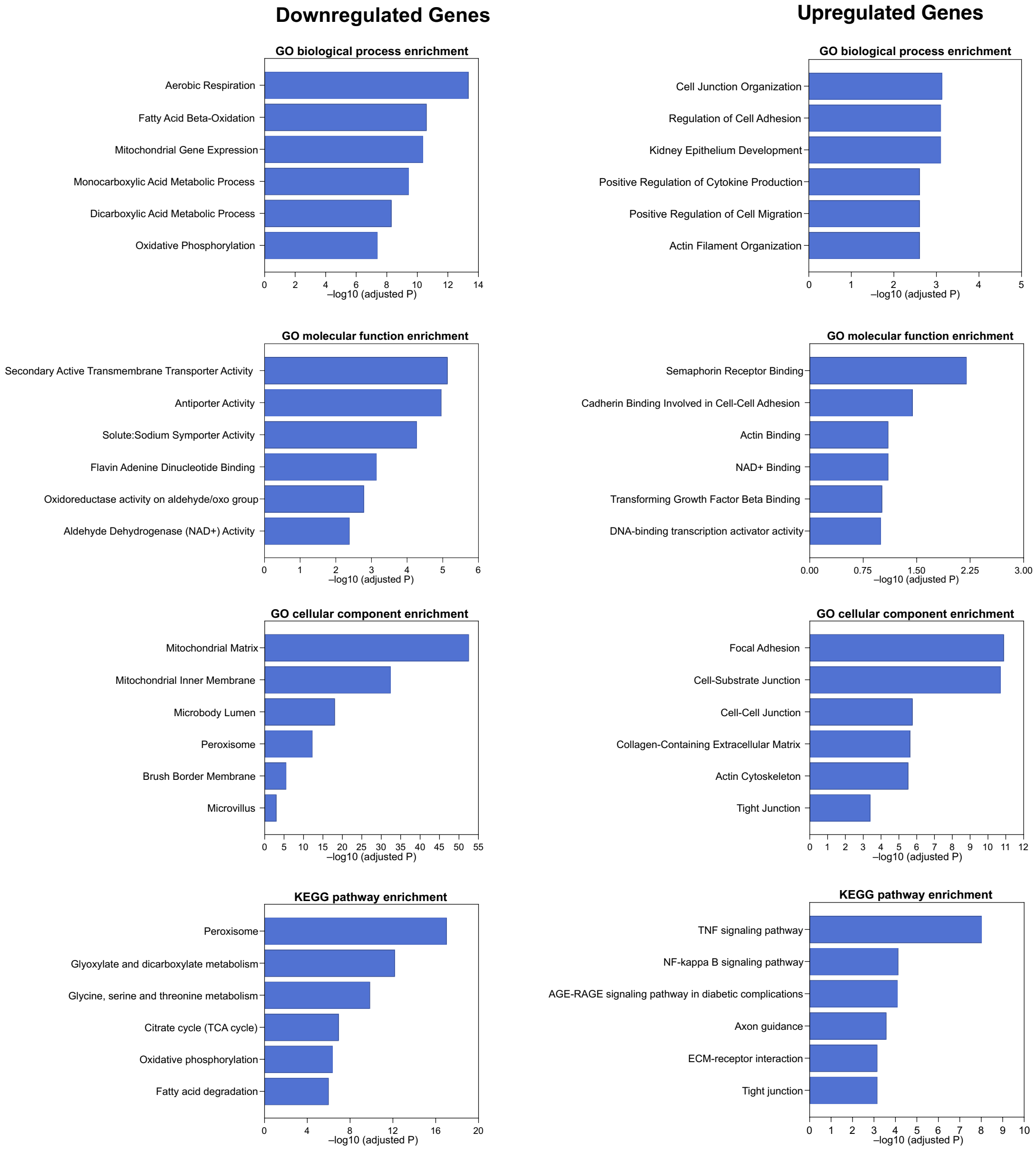


**Supplemental Figure 3. Gene Ontology and KEGG pathway enrichment analyses of candidate direct HNF4A target genes identified by integrated RNA-seq and CUT&RUN.** GO biological process, molecular function, and cellular component and KEGG pathway enrichment analyses were performed for candidate direct HNF4A target genes that were downregulated (left panels) or upregulated (right panels) following *Hnf4a* deletion. Downregulated targets were enriched for transport and metabolic pathways, consistent with loss of proximal tubule functional programs. In contrast, upregulated targets were enriched for pathways related to cell adhesion, extracellular matrix organization, and epithelial remodeling. Bars represent the top enriched terms ranked by significance.

Peaks associated with downregulated genes:

| Rank | Motif | *P* value | % Target | TF |
|---|---|---|---|---|
| 1 | 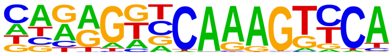 | 1e-1836 | 44.47% | HNF4A |
| 2 | 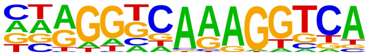 | 1e-927 | 44.17% | PPAR |
| 3 | 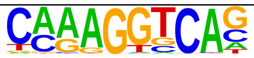 | 1e-547 | 56.82% | ESRRA |
| 4 | 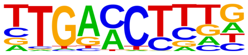 | 1e-540 | 63.54% | RARA |
| 5 | 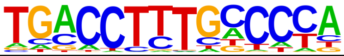 | 1e-503 | 32.53% | PPARE |
| 6 | 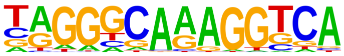 | 1e-490 | 35.54% | RXR |
| 7 | 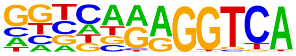 | 1e-356 | 37.03% | COUP-TFII |
| 8 | 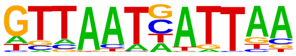 | 1e-337 | 7.80% | HNF1B |

Peaks associated with upregulated genes:

| Rank | Motif | *P* value | % Target | TF |
|---|---|---|---|---|
| 1 | 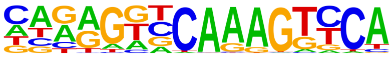 | 1e-1056 | 41.78% | HNF4A |
| 2 | 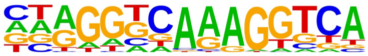 | 1e-513 | 41.71% | PPAR |
| 3 | 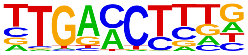 | 1e-308 | 60.97% | RARA |
| 4 | 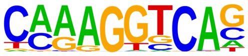 | 1e-302 | 54.85% | ESRRA |
| 5 | 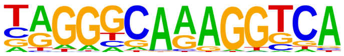 | 1e-275 | 34.23% | RXR |
| 6 | 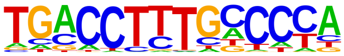 | 1e-271 | 30.76% | PPARE |
| 7 | 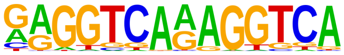 | 1e-214 | 9.60% | TR4 |
| 8 | 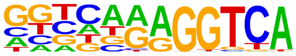 | 1e-198 | 35.26% | COUP-TFII |
| 9 | 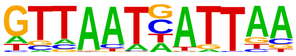 | 1e-154 | 6.32% | HNF1B |

**Supplemental Figure 4. HNF4A CUT&RUN peaks associated with differentially expressed genes are enriched for HNF4A and other transcription factor motifs.** Motif enrichment analysis of HNF4A CUT&RUN peaks associated with genes downregulated (top) or upregulated (bottom) following *Hnf4a* deletion. The canonical HNF4A motif is the most significantly enriched motif in both datasets. Motifs recognized by PPAR, ESRRA, RAR, RXR, COUP-TFII, and HNF1B are also significantly enriched, suggesting potential co-regulatory interactions. Tables show motif rank, statistical significance (*P* value), percentage of target regions containing each motif, and corresponding transcription factor.

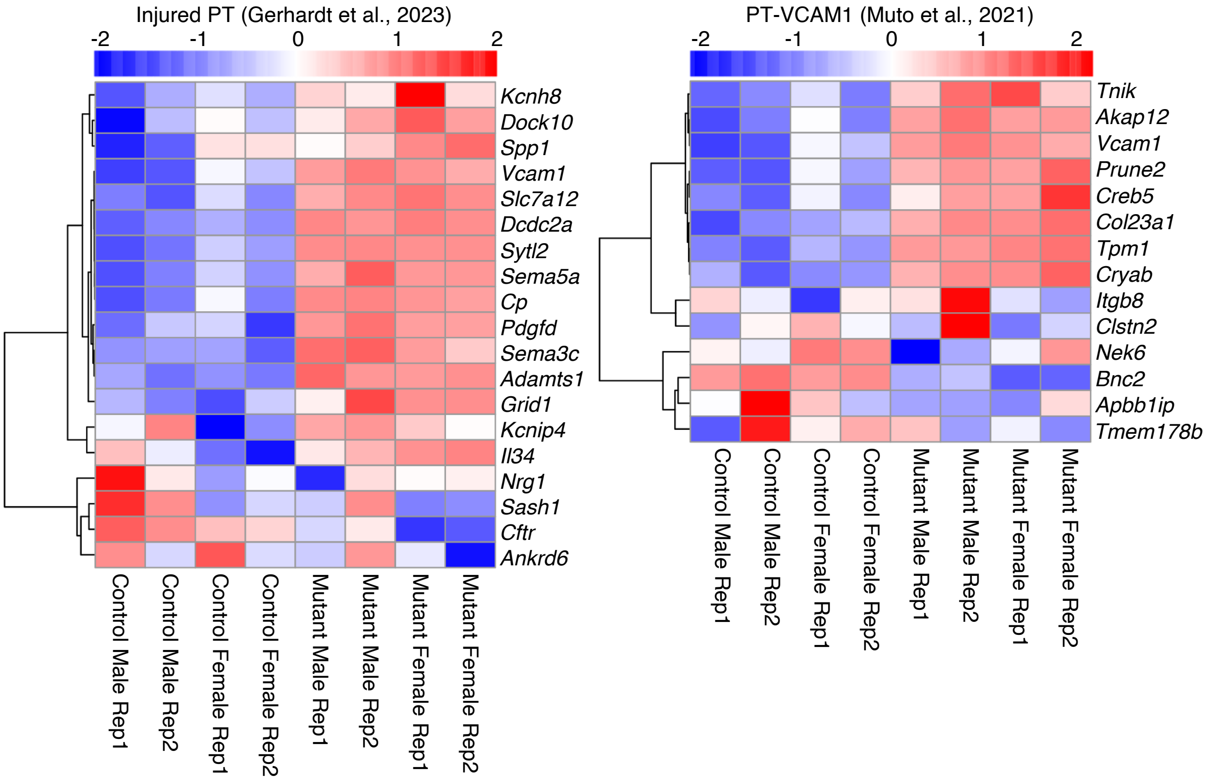


universe size=19141
set1_size=2182
set2_size=19
overlap=10
expected=2.1659
fold enrichment=4.6170
$P = 1.3 \times 10^{-5}$

universe size=19141
set1_size=2182
set2_size=14
overlap=5
expected=1.5959
fold enrichment=3.1329
$P = 1.6 \times 10^{-2}$

**Supplemental Figure 5. *Hnf4a* mutant proximal tubules show enrichment of the Injured PT and PT-VCAM1 gene signatures.** Heatmaps showing expression of Injured PT signature genes identified by Gerhardt et al. (left) and PT-VCAM1 signature genes described by Muto and Wilson et al. (right) in bulk RNA-seq datasets from control and *Hnf4a* mutant proximal tubules. Gene expression values are displayed as row-scaled Z-scores. Compared with control samples, *Hnf4a* mutant proximal tubules exhibit coordinated upregulation of genes associated with Injured PT and the PT-VCAM1 transcriptional program. Sample sex and genotype are indicated below each heatmap.

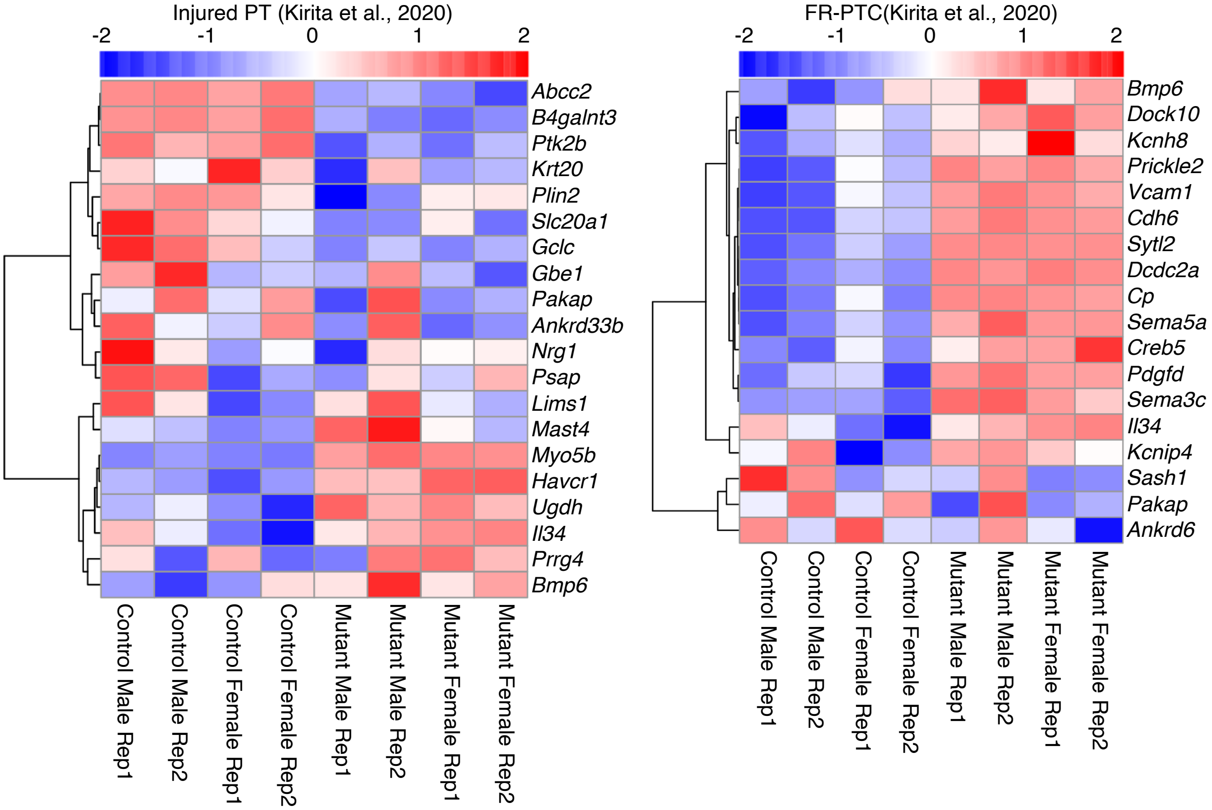


universe size=19141
set1_size=2182
set2_size=20
overlap=3
expected=2.2799
fold enrichment=1.3158
$P = 4.0 \times 10^{-1}$

universe size=19141
set1_size=2182
set2_size=18
overlap=8
expected=2.0519
fold enrichment=3.8988
$P = 4.3 \times 10^{-4}$

**Supplemental Figure 6. *Hnf4a* mutant proximal tubules show preferential upregulation of the FR-PTC gene signature compared with the injured PT signature.**
Heatmaps showing expression of injured PT (left) and FR-PTC (right) signature genes defined by Kirita et al. in RNA-seq datasets from control and Hnf4a mutant proximal tubules. Gene expression values are displayed as row-scaled Z-scores. Compared with control samples, Hnf4a mutant proximal tubules show limited changes in the injured PT gene signature but prominent upregulation of the FR-PTC signature, consistent with acquisition of a failed repair-associated proximal tubule transcriptional program. Sample sex and genotype are indicated below each heatmap.

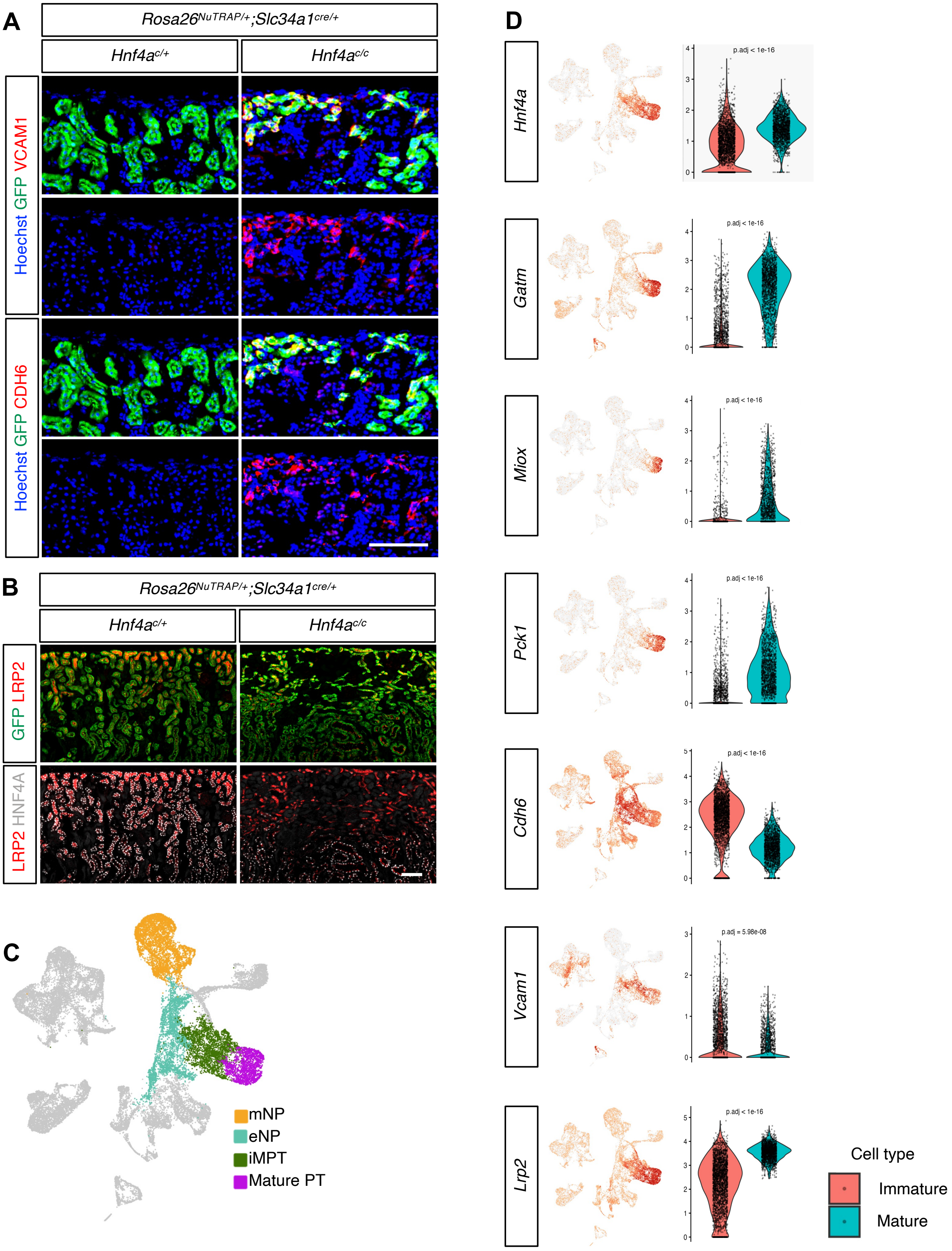
A
Rosa26NuTRAP/+;Slc34a1cre/+
Hnf4ac/+
Hnf4ac/c
Hoechst GFP VCAM1
Hoechst GFP CDH6
B
Rosa26NuTRAP/+;Slc34a1cre/+
Hnf4ac/+
Hnf4ac/c
GFP LRP2
LRP2 HNF4A
C
mNP
eNP
iMPT
Mature PT
D
Hnf4a
p.adj < 1e-16
Gatm
p.adj < 1e-16
Miox
p.adj < 1e-16
Pck1
p.adj < 1e-16
Cdh6
p.adj < 1e-16
Vcam1
p.adj = 5.98e-08
Lrp2
p.adj < 1e-16
Cell type
Immature
Mature

**Supplemental Figure 7. Loss of HNF4A is associated with acquisition of an immature proximal tubule–like state.** (A) Co-immunostaining for GFP, VCAM1, and CDH6 in control and *Hnf4a* mutant kidneys demonstrates co-detection of VCAM1 and CDH6 in GFP+/HNF4A-deficient proximal tubule cells. Stage: P61; scale bar: 100 µm. (B) Immunofluorescence staining for LRP2 in control and *Hnf4a* mutant kidneys demonstrates retention of LRP2 in a subset of GFP+/HNF4A-deficient proximal tubules. Stage: P61; scale bar: 100 µm. (C) UMAP visualization of the reference single-cell RNA-seq dataset showing mesenchymal nephron progenitors (mNP), epithelial nephron progenitors (eNP), immature proximal tubules (iMPT), and mature proximal tubules (Mature PT). (D) Feature plots and corresponding violin plots showing expression of *Hnf4a, Gatm, Miox, Pck1, Cdh6, Vcam1,* and *Lrp2* across immature and mature proximal tubule populations. *Hnf4a* and *Lrp2* are expressed in both immature and mature proximal tubules but at higher levels in mature proximal tubules. *Gatm, Miox,* and *Pck1* are predominantly expressed in mature proximal tubules, whereas *Cdh6* and *Vcam1* are enriched in immature proximal tubule populations. **Abbreviations:** mNP, mesenchymal nephron progenitor; eNP, epithelial nephron progenitor; iMPT, immature proximal tubule; Mature PT, mature proximal tubule.

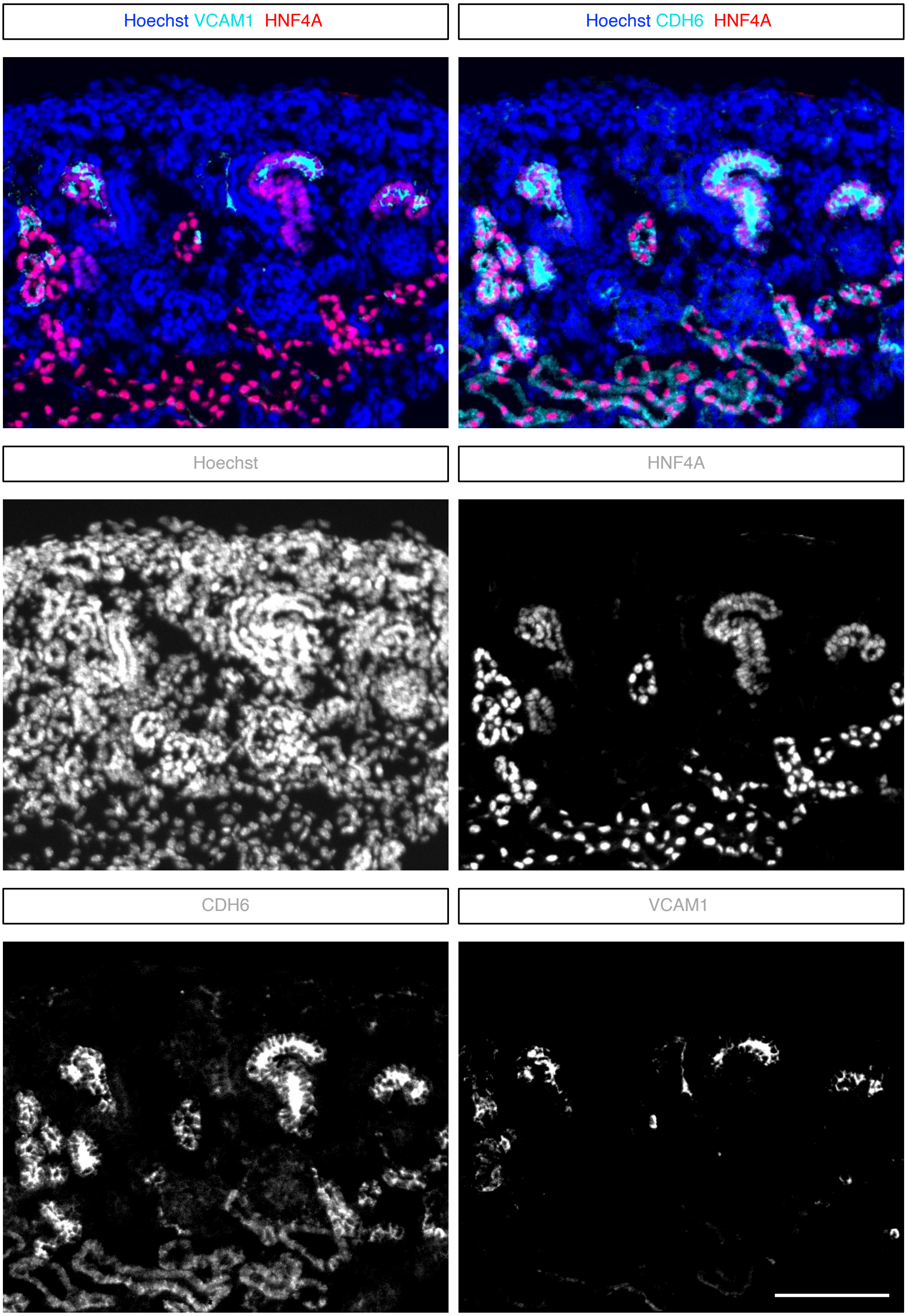


**Supplemental Figure 8. VCAM1 is detected in a subset of CDH6-high proximal tubule progenitor cells in the developing kidney.** VCAM1 is detected in a subset of CDH6-high proximal tubule progenitor cells, showing partial overlap of these markers during nephrogenesis. Stage: E18.5; scale bar: 100 µm.

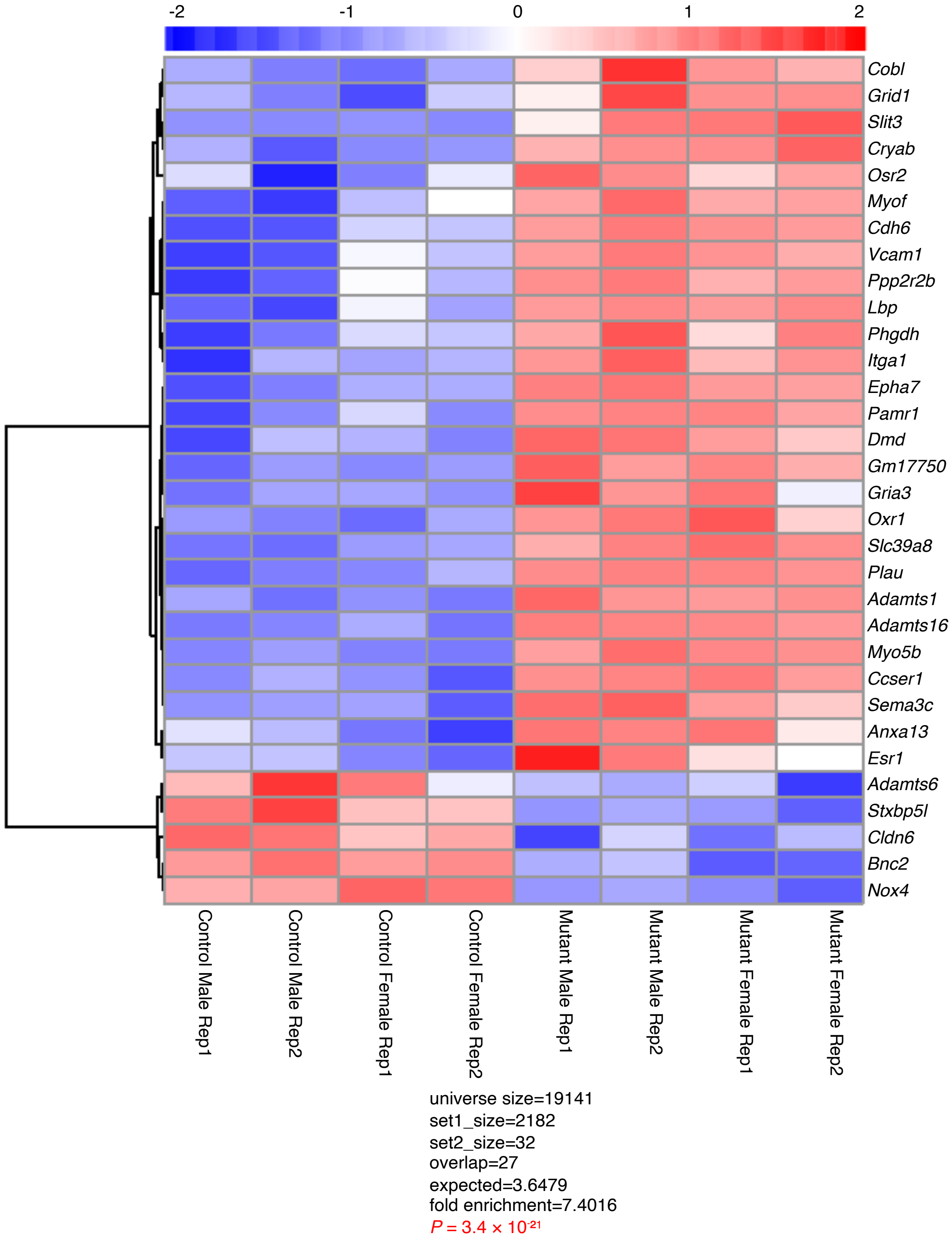


**Supplemental Figure 9. HNF4A-deficient proximal tubules acquire an immature proximal tubule–like transcriptional state.** Heatmap showing expression of 32 differentially expressed immature proximal tubule–associated genes in RNA-seq samples from control and *Hnf4a* mutant proximal tubules. Gene expression values are displayed as row-scaled Z-scores. Compared with control samples, *Hnf4a* mutant proximal tubules show predominant upregulation of immature proximal tubule–associated genes, consistent with acquisition of an immature proximal tubule–like transcriptional state. Sample sex, genotype, and replicate are indicated below the heatmap.
